# hvantk: an agentic multi-omics framework for variant annotation and analysis

**DOI:** 10.64898/2026.10.19.760196

**Authors:** Enrique Audain, Yasset Perez-Riverol, Marc-Phillip Hitz

## Abstract

Large-scale genetic variant analysis involves multiple computational steps, data formats, gene and variant annotation sources and compute requirements. Some of the steps like joint calling for large cohorts are run on large compute clusters, while gene/variant classification algorithms can be run in a desktop computer. At the same time, variant annotations such as gene expression and gene constraints metrics from gnomAD are provided in different formats, by resources that frequently change their versions. Here, we present *hvantk*, a framework that provides a common data model for large-scale annotation and analysis of genetic variants, by extending popular data formats such as Hail MatrixTable and AnnData. hvantk agentically integrates 21 public multi-omic data providers as plugins that a developer, bioinformatician or an AI agent can author without modifying the framework’s core, drives analysis workflows using agent skills, and scales from single desktop computers to high-performance computing clusters. We used hvantk for reanalysing three case-control cohorts, comprising congenital heart disease (CHD), developmental and epileptic encephalopathy (Epi25) and schizophrenia (SCHEMA). Re-ranking of gene-based burden testing results using multi-omic annotations improved the recovery of curated disease genes from 26% to 49% in CHD, 40% to 48% in Epi25 and 51% to 54% in SCHEMA, among the top decile re-ranked genes. Expression at bulk and single-cell levels were the features contributing the most over a baseline of burden, gene constraint and publication count.

## Background

Scalable genetic variant analysis is a multi-stage computational process that starts from raw sequence reads through per-sample variant calling, cohort joint genotyping and multi-omics annotation to interpretation. Each of these steps involves multiple tools, file formats, and data analysis workflows. For example, sarek [1] takes raw sequencing data to per-sample variant call format (VCF) files, whereas Hail [2] is used to develop pipelines that allow further analysis of cohort-level data. In addition, depending on the dataset and the analysis step, data must be processed in different computational environments, from single desktop computers to large-scale distributed processing. The analysis therefore moves through stages that demand distinct computational resources, changes data formats, and algorithms that need different inputs or execution environments. These challenges can be grouped into three major issues: (i) provenance, (ii) file format heterogeneity, and (iii) scalability.

As annotation sources and databases evolve, new releases can reclassify variants or revise gene annotations such as constraint metrics [3]. Provenance and source-version tracking are therefore recognised requirements for FAIR analysis [4], yet they remain poorly supported by many annotation tools and are not broadly adopted by analytic pipelines. Recent initiatives such as BioClaw [5] include expert-curated workflows into versioned agent skills, but do not track changes in the upstream data sources on which those skills depend. Genetic variant and gene analysis also involves heterogeneous data representations. A few examples include genomic VCF (gVCF) generated with tools such as GATK [6], Hail *MatrixTables* for cohort-level analysis [2], and AnnData objects for gene-level expression data integration [7]. These stages differ not only in data structure but also in computational requirements, ranging from distributed processing to analysis on single desktop computers. Interoperable data models that preserve provenance across data sources and execution environments are therefore important when combining cohort-level association results with external biological evidence to improve the interpretation of disease-relevant variants and genes [8–10].

These challenges are widely represented in population genomics association studies. Existing tools for association analysis [11,12], that link variants or genes to a specific disease or phenotype, provide the primary statistical evidence. However, selecting candidate variants and genes for biological interpretation or follow-up often requires additional context. For example, gene expression in a disease-relevant tissue can provide evidence that a statistically associated gene is also biologically plausible [13]. Integrating further biological evidence, such as variant-level prediction scores, bulk or single-cell expression and post-translational modifications, requires compatible data representations, traceable annotations, and methods that combine cohort-level association results with external biological information.

Here we present hvantk (https://github.com/bigbio/hvantk), a multi-omic agentic framework for large-scale variant annotation and analysis. The framework implements a common data model that extends Hail’s and AnnData file formats, which facilitate translation and serialization between data types and formats, and increase interoperability of its core algorithms and pipelines. On top of these core data structures, hvantk retrieves annotations from public multi-omics resources such as Expression Atlas [14] and PeptideAtlas [15] through data plugins. An AI coding agent can update a data provider or add a new one, and can drive analysis workflows through skills. The resulting data are converted into hvantk’s data structures, with version and provenance information recorded and source-specific checks for upstream changes, allowing integration with *in-house* variant data. The analytic runtime itself remains deterministic and independent of large language models (LLMs). hvantk allows for the core algorithms to run in specific execution environments depending on the data scale, from Spark or high-performance computing (HPC) clusters for cohort-level processing to local execution for aggregated tabular data analyses. We integrate 21 public resources and apply hvantk to per-sample gVCFs from an *in-house* congenital heart disease (CHD) case-control exome cohort, combining burden statistics with biological annotations to generate a ranked per-gene score. We extended the analysis by using aggregated burden statistics from two additional case-control disease cohorts, the developmental and epileptic encephalopathy (Epi25) and schizophrenia (SCHEMA) cohorts. Across the three cohorts, we measure what multi-omics annotations add to the ranking. Single-cell and bulk multi-organ expression explained most of the signal gained over a baseline model, whereas eQTL and PTM features showed negligible contribution.

## Results

### The hvantk framework

The hvantk framework consists of four major components: (i) a core data model and serialization layer; (ii) agent-extensible multi-omics data-provider plugins; (iii) analysis tools and algorithms; and (iv) an AI-assisted layer comprising plugin and workflow skills that guide AI coding agents (**Figure 1**). Data can flow across pre-built pipelines, including joint genotyping, quality control (QC), burden testing, annotation across 21 multi-omics providers and the application of algorithms and tools. The framework is multi-backend: a common data model extends Hail *MatrixTable* for variant- and cohort-scale data [2], AnnData [7] for expression data, and pandas for *in-memory* tabular data analyses. Distributed computing is therefore used only for stages that require it.

**Figure 1.**
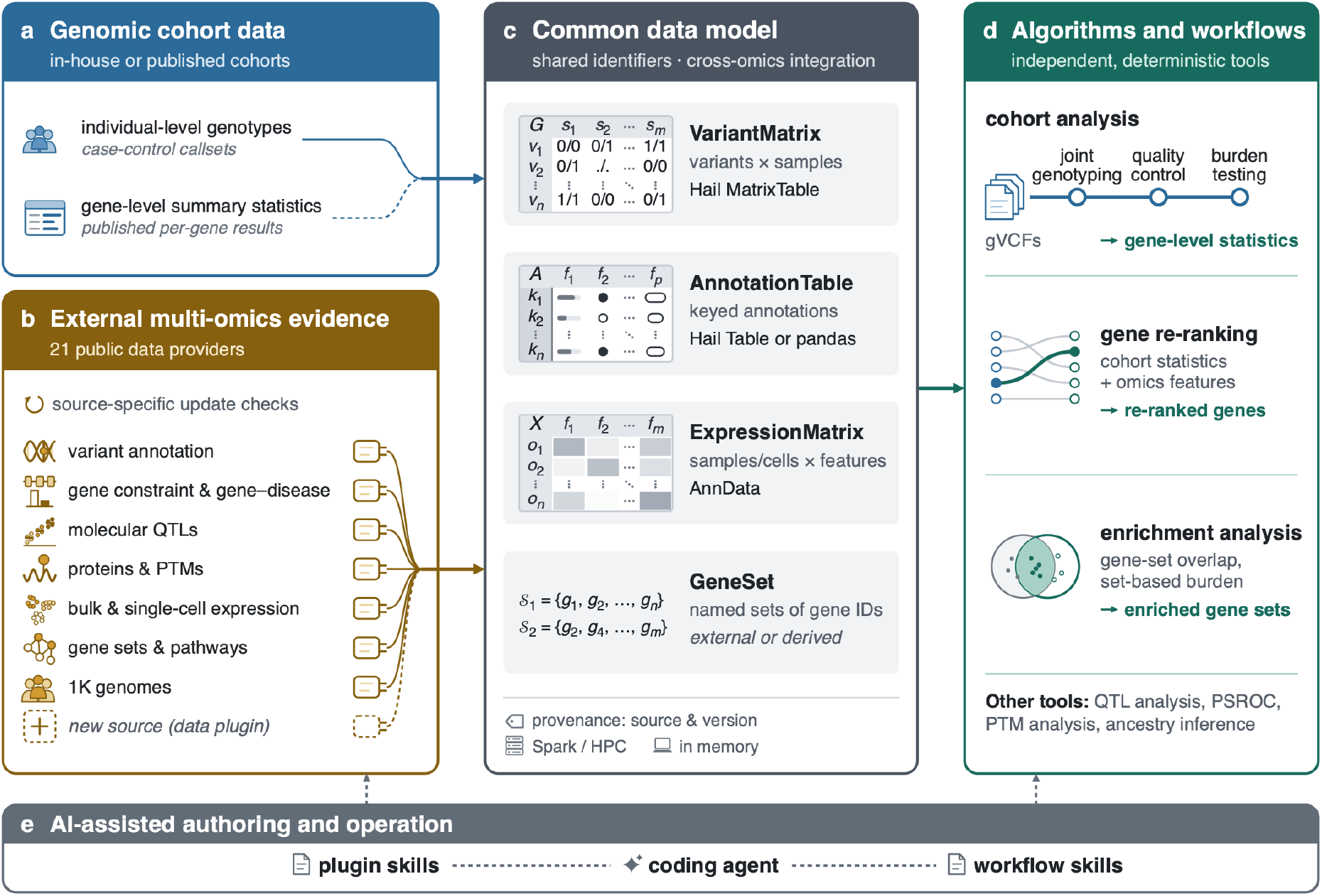
Overview of the *hvantk* framework for genetic variants multi-omic annotation and analysis. **(a)** Genomic cohort data can be provided to the framework in two ways: individual-level genotype data, or alternatively, as previously computed gene-level (dashed line) burden or association statistics. **(b)** External biological resources are integrated through data plugins from 21 public providers (**Table 1**). Plugins support source-specific update checks and can be developed or updated with AI coding agents. **(c)** A common data model represents these resources as four data types (VariantMatrix, AnnotationTable, ExpressionMatrix and GeneSet), with shared identifiers and provenance records supporting cross-omics integration. Processing can use distributed (Spark/HPC clusters) or *in-memory* computation according to data scale and analysis requirements. **(d)** Datasets support independent algorithms and workflows, including joint genotyping and quality control of per-sample gVCFs with HGC (Figure 2), gene burden testing, gene re-ranking, enrichment analysis and QTL analysis. **(e)** AI-assisted data plugin development and workflow orchestration. Plugin skills guide a coding agent in developing data plugins. Workflow skills guide the agent in orchestrating analysis workflows. The underlying tools execute deterministically and independently of LLMs.

**Table 1.** Public data providers integrated by hvantk as plugins. . The 21 providers are implemented as independent data plugins, some of which generate multiple datasets (e.g. collection-like providers such as UCSC Cell Browser). Each dataset carries a provenance record. Providers are grouped by biological data type and primary identifiers. Access indicates whether hvantk provides a built-in downloader or instructions for obtaining the source data.

| Provider | Dataset type | Access | Reference |
| --- | --- | --- | --- |
| *Variants* (keyed by locus, alleles) |  |  |  |
| ClinVar | *AnnotationTable*: clinical significance | downloader | [18] |
| dbNSFP | *AnnotationTable*: pathogenicity scores | documented | [19] |
| AlphaGenome | *AnnotationTable*: variant effect predictions | documented | [20] |
| 1000 Genomes | *VariantMatrix* + samples | downloader | [17] |
| GWAS Catalog | *AnnotationTable*: GWAS trait associations | downloader | [21] |
| *Gene constraint / gene-disease* (keyed by gene_id or hgnc_id) |  |  |  |
| gnomAD metrics | *AnnotationTable*: gene constraint | downloader | [22] |
| GeVIR | *AnnotationTable*: gene constraint | downloader | [23] |
| ClinGen | *AnnotationTable*: gene-disease validity | downloader | [24] |
| GenCC | *AnnotationTable*: gene-disease curation | downloader | [25] |
| COSMIC CGC | *AnnotationTable*: cancer gene census | documented | [26] |
| *Molecular QTLs* (keyed by locus, alleles, gene_id) |  |  |  |
| GTEx eQTL | *AnnotationTable*: eQTL summary stats | documented | [27] |
| pQTL | *AnnotationTable*: pQTL summary stats | documented | [28] |
| *Proteins* (keyed by genomic interval / protein position) |  |  |  |
| INSIDER | *AnnotationTable*: protein interactions | downloader | [29] |
| UniProt PTM | *AnnotationTable*: PTM sites | downloader | [30] |
| PeptideAtlas | *AnnotationTable*: phosphoproteome | downloader | [15] |
| *Expression / abundance* (feature × sample matrices; feature = gene or PTM site) |  |  |  |
| Expression Atlas | *ExpressionMatrix*: bulk RNA-seq | downloader | [14] |
| UCSC Cell Browser | *ExpressionMatrix*: scRNA-seq | downloader | [31] |
| CPTAC | *ExpressionMatrix*: tumor proteome / phosphosite intensity | downloader | [32] |
| *Gene sets / pathways* (keyed by set_name) |  |  |  |
| MSigDB | *AnnotationTable*: gene-set / pathway definitions | documented | [33] |
| *Mapping / reference* (keyed by gene_id / hgnc_id) |  |  |  |
| Ensembl | *AnnotationTable*: gene structure | downloader | [34] |
| HGNC | *AnnotationTable*: gene nomenclature | downloader | [35] |

Datasets built with hvantk can be grouped into four types: *AnnotationTable*, *ExpressionMatrix*, *VariantMatrix*, or *GeneSet*. Each data structure records its source plugin, version, upstream fingerprint, and parent provenance, allowing derived datasets and cross-omics joins to be traced back to their contributing sources and versions. The implemented data-plugin system makes hvantk AI agent-extensible. Each provider is a self-contained plugin defined by a schema-validated specification and new sources can be added without modifying the framework codebase. When an AI coding agent adds a provider, it must work against validated schemas rather than inferred ones. Unlike broader definitions of agentic genomics, in which autonomous LLM-based agents execute multi-step analyses [16], hvantk uses dedicated skills to guide AI coding agents in developing data plugins and orchestrating analysis workflows. The following sections show how these capabilities are applied to analyse case-control exome cohorts.

### Large-scale genome analysis: joint genotyping with HGC

hvantk includes the Hail Genotype Combiner (HGC), a five-stage pipeline that takes per-sample gVCFs and assembles them into an analysis-ready cohort callset (**Figure 2a**). It does not re-implement joint genotyping: the combine step uses the Hail combiner, and scalability rests on the Hail Scalable Variant Call Representation (SVCR) [2]. hvantk contributes the orchestration instead: one command runs combine, incremental merge-and-append, densification, multi-sample VCF export with allele counts recomputed on the joint callset, and quality control. Because of the SVCR architecture, new samples can be added to the callset without re-genotyping the existing cohort.

**Figure 2.**
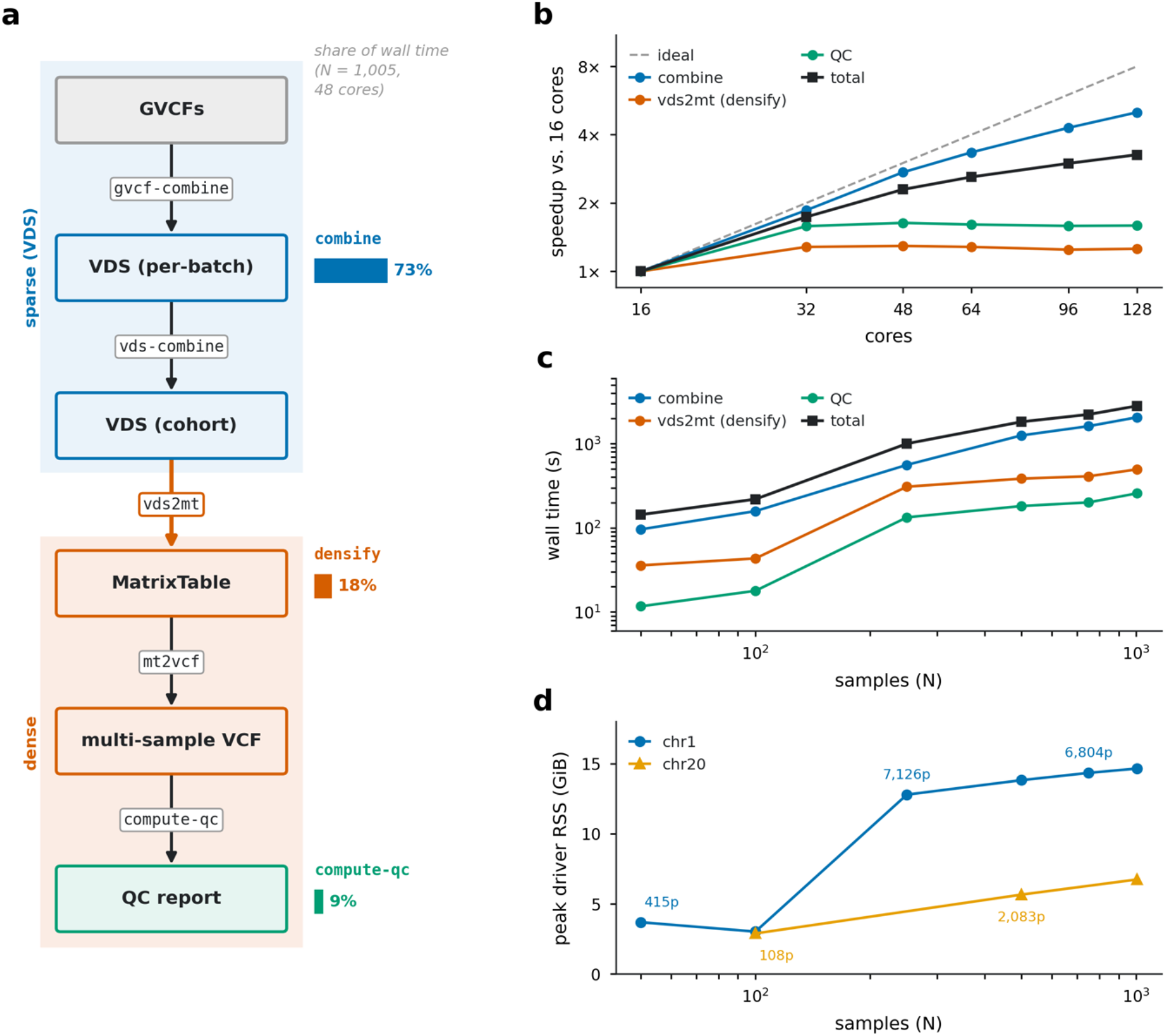
Cohort-scale joint genotyping with HGC. Per-sample whole-genome sequencing gVCFs from a fraction (n=1,005) of the primary CHD cohort were used for a demonstrative scalability test on a Spark cluster. **(a)** The five stages, run by one *hvantk hgc* pipeline sub-command with resumable per-stage checkpoints. Shading separates the sparse (VDS) and dense matrix representations, and bars denote each stage’s share of wall time at n=1,005 on 48 cores. **(b)** Scaling at N=500 over 16-128 cores, as speedup over the same stage’s 16-core wall time. **(c)** Cohort scaling: per-stage wall time against N at a fixed 48 cores. **(d)** Peak driver resident memory against N, for chr1 and chr20, with selected points labelled with their VDS partition count. Quality control metrics of the joint callset (1,005 samples and 3,928,030 polymorphic variants) are shown in **Additional file 1: Fig. S2.**

A fraction (n=1,005) of the primary CHD cohort [10] was used for a demonstrative scalability test (**Additional file 1: Fig. S1**). We tested how runtime varies with the number of CPUs in a Spark cluster for a fixed number of samples (**Figure 2b**). The combine step dominates the compute cost and continues to scale, reaching a 5-fold speedup at 128 cores, whereas densification and quality control saturate at 32 cores. The reason is that the partition layout of the native Hail Variant Dataset (VDS) format follows the cohort’s reference-block count, not the size of the dense matrix that is written. When the cohort size is varied on a fixed Spark cluster, the number of reference blocks saturates once every region of the genome is already represented by a block in some sample; adding further samples therefore creates no new blocks, whereas the dense matrix keeps growing with N. The densification step costs 2.9-fold more per sample at N=250 than at N=100 (**Figure 2c**). Peak driver memory likewise tracks partition count, not sample count. The same 1,005 samples need 6.7 GiB with 1,941 partitions but 14.7 GiB with 7,035 partitions (**Figure 2d**).

hvantk includes other capabilities which are standard when assembling a cohort: the computation of per-variant and per-sample quality-control metrics (**Additional file 1: Fig. S2**) and sample ancestry inference using reference population datasets (e.g. the 1000 Genomes Project (1KG) [17]). The primary case-control cohort (CHD) used in the present study was generated using these methods, established in a companion study [10]. A callset is then passed to the downstream stages, starting with annotation, which is handled by the data plugin module of hvantk.

### Agent-extensible multi-omics annotation across data providers

#### Annotation sources and providers

hvantk integrates 21 public data providers as self-contained plugins across five biological levels: genetic variants, genes and gene-disease relations, molecular QTLs, protein features, and expression (**Table 1**). Because the providers resolve to common data models, a join operation is possible without extra manual schema reconciliation. Fifteen of the 21 providers include a built-in downloader, covering 16 of the 26 datasets provided with the framework. The rest are too large or require a licence to fetch automatically, and therefore, how to get the primary data is documented instead. UCSC Cell Browser [31] and Expression Atlas [14] are collection-like resources, exposing a builder parameterized by accession over committed lists of 270 single-cell and 27 bulk RNA-seq accessions. Expression features can therefore be matched to a disease-specific tissue by selecting a dataset rather than by adding a new data plugin.

#### Extending the annotation framework using AI agents

New annotation resources can be integrated into hvantk by an AI coding agent without modifying the core library (**Figure 3a**). For each resource, the agent implements the data processing workflow (builder) and validates the resulting dataset against predefined schemas and reference data snapshots. Once validated, plugins generate datasets through deterministic execution, which can be combined into a feature matrix (**Figure 3b**). The framework includes automated tests to detect changes in both dataset generation and upstream annotation sources. Because external databases evolve over time, potentially changing variant classifications (e.g. ClinVar) and/or gene-level metrics [3,36], source-specific checks monitor these changes and generate draft updates for review before incorporation into downstream analyses. This process depends on source availability: sources like gnomAD constraint tables are monitored using file checksums and size, whereas dbNSFP is checked for new releases. Some data providers have licence restrictions, and synthetic test data are used instead where the original data cannot be redistributed (**Additional file 1: Fig. S3**).

**Figure 3.**
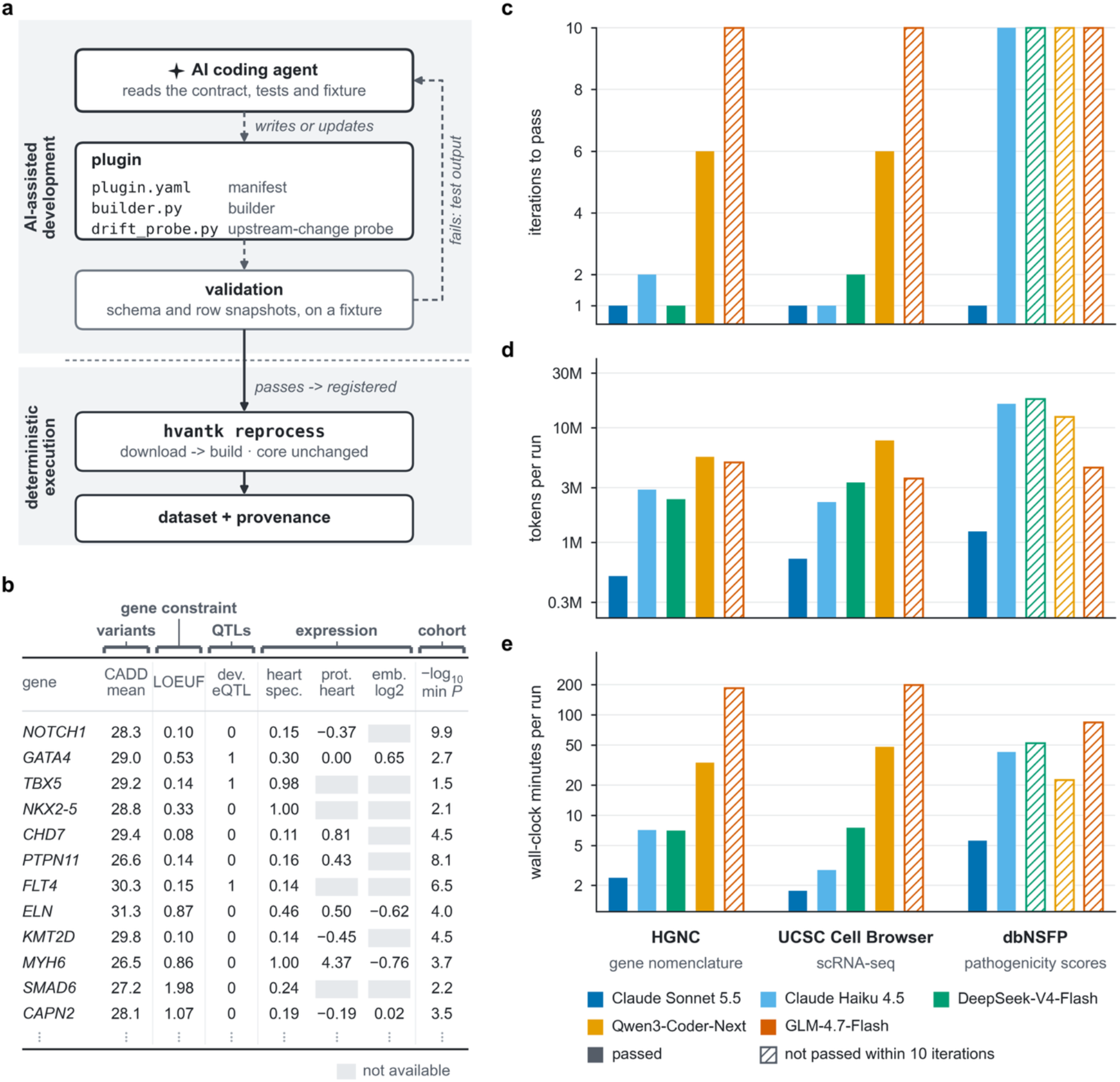
AI-assisted data plugin integration. **(a)** Extending the annotation module and validation workflow. An AI coding agent creates or updates a data plugin against its specification. The agent iteratively validates its implementation against reference schemas and data snapshots, using test failures as feedback. Once validated, the plugin is executed by hvantk to generate datasets with recorded provenance. **(b)** Example of a gene-level feature matrix assembled from standardized annotation datasets and cohort-derived statistics, joined by shared gene identifiers. The table shows a representation of the CHD dataset. (c-e) Benchmark of five AI coding models developing plugins for three annotation providers (HGNC, UCSC Cell Browser and dbNSFP), with three independent runs per model-provider combination and a maximum of ten iterations. **(c)** Iterations required to pass unit tests. **(d)** Token consumption per run. **(e)** Wall-clock time per run. Bars represent the median across three runs, with colors identifying AI coding models. Hatched bars indicate combinations in which the median run did not pass the tests within ten iterations. Token consumption and wall-clock time are shown on logarithmic scales.

We compared five AI coding models across three data providers (HGNC, UCSC Cell Browser and dbNSFP) to evaluate success rate on developing a specific data plugin (**Figure 3c-e**). Claude Sonnet 5.5 passed the unit tests in all performed runs, whereas success rate varied across other models such as Claude Haiku 4.5, DeepSeek-V4-Flash, Qwen3-Coder-Next and GLM-4.7-Flash. Token consumption and execution time also varied considerably depending on the model and provider, indicating differences in both the models’ reliability and computational cost.

#### Relation to existing annotation tools

hvantk is not intended to replace established annotation tools that already assemble per-variant and -gene evidence across many providers [37]. hvantk addresses a different property of the annotation layer itself. Every dataset carries provenance and an upstream-change check, and new sources are authored against a validated schema that an AI agent can satisfy without modifying the codebase. The resulting datasets can be used directly by the algorithms and methods included in the library. A given analysis runs on a configured subset of a data catalogue, as shown below, where gene-based burden statistics from case-control exome studies are re-ranked using multi-omics features assembled from different data providers.

### Re-ranking per gene burden statistics using multi-omics features

We evaluate the framework by asking whether multi-omics evidence improves the interpretation of a burden result beyond the gene-prior scores [23,38–40] that an analyst routinely uses. We analysed three cohorts: the developmental and epileptic encephalopathy collection (Epi25) [8], the SCHEMA schizophrenia meta-analysis [9], and a congenital heart disease (CHD) cohort [10]. CHD is the study primary cohort, since it is run through the full toolkit and supplies the callset built by the HGC pipeline described previously. However, the re-ranking analysis was performed for all three cohorts. Four experimental design aspects are held fixed for each cohort: i) every gene with a burden result was included, without conditioning on the burden statistic, ii) positive labels used during training are defined based on expert curation and sources like ClinGen [24] and GenCC [25], iii) a fixed baseline against which every increment is measured, including the burden statistic, gene constraint metrics and publication count, and iv) the gene-prior scores against which the re-ranking score is compared (**Methods**). **Table 2** defines the terms used throughout.

**Table 2.** Glossary of terms.

| Term | Description |
| --- | --- |
| *SVCR and Variant Dataset (VDS)* | The Scalable Variant Call Representation (SVCR) is Hail's sparse representation of cohort sequencing data, implemented as a Variant Dataset (VDS) containing separate reference-block and variant-data components. |
| *Hail MatrixTable* | A distributed matrix-like data structure for cohort-scale genomic data, with variants as rows, samples as columns and genotype-level information as entries. |
| *AnnData* | An annotated matrix format widely used for single-cell data; hvantk uses AnnData to represent expression matrices and associated metadata. |
| *Gene-prior score (DOMINO, shet, pHaplo, GeVIR)* | Published gene-level scores that estimate how likely a gene is to be involved in disease, derived from population and evolutionary data and computed independently of any study cohort. |
| *Post-translational modification (PTM) density* | The number of UniProt-annotated protein modification sites (e.g. phosphorylation) mapped to the codons of a gene. |
| *Positive-unlabelled (PU) setting* | A label setting in which confirmed disease genes form the positive class, while genes without a curated association remain unlabelled and may include undiscovered disease genes. |
| *ROC-AUC and Average Precision (AP)* | Measures of discrimination. ROC-AUC quantifies ranking of positives relative to negatives, whereas average precision summarises precision across recall and is informative under class imbalance. |
| *Top-decile recall and precision* | Top-decile recall is the share of curated disease genes ranked in the top 10% of the gene universe. Precision among the top 100 genes is the share of those 100 genes that are curated disease genes. |
| *Out-of-fold prediction and group cross-validation* | A score is out of fold when it comes from a model that did not see that gene during training. In group cross-validation, genes from the same HGNC gene group are kept in the same fold, so that a gene is never scored by a model trained on a related gene. |
| *Paired bootstrap confidence interval* | A confidence interval obtained by resampling genes with replacement while preserving the same resample across models, allowing uncertainty in model differences to be estimated on paired data. |
| *Provenance record and upstream fingerprint* | A provenance record stores the source, version, build time and parent datasets used to generate a dataset. An upstream fingerprint records the state of the source and is used to detect subsequent changes. |
| *Data plugin* | A data plugin defines how an external resource is retrieved and converted into hvantk data structures, through a schema-validated specification and a source-specific check for upstream changes. |
| *Snapshot test* | A regression test comparing a rebuilt dataset with a stored reference schema and representative rows to detect unexpected changes. |

#### Omics evidence improves ranking in all three cohorts

Added to the baseline, omics features improve gene ranking in every cohort: ΔAUC +0.089 in CHD, +0.054 in Epi25 and +0.028 in SCHEMA, all confidence intervals excluding zero. Added instead to a model that includes the gene prior scores fitted, the margins are still larger (+0.139, +0.083 and +0.031), so the result does not depend on whether the priors are refitted or used as originally published **(Additional file 1: Tables S1** and **S2**). **Figure 4a** compares three ranking scores, the published gene priors, the baseline, and the baseline extended with multi-omics evidence, by counting how many curated disease genes each places in the top 10% of the cohort gene universe. In the evaluated CHD cohort, curated disease genes among the top 100 ranked genes rise from 5% to 39% (precision). The choice of comparator also matters: AUC achieved by the four published priors ranges from 0.622 to 0.670 individually and 0.646 in combination, whereas publication count alone reaches 0.663. Therefore, how often a gene is mentioned in publications, a count that grows as genes are studied, ranks CHD genes about as well as the evaluated gene prior scores.

**Figure 4.**
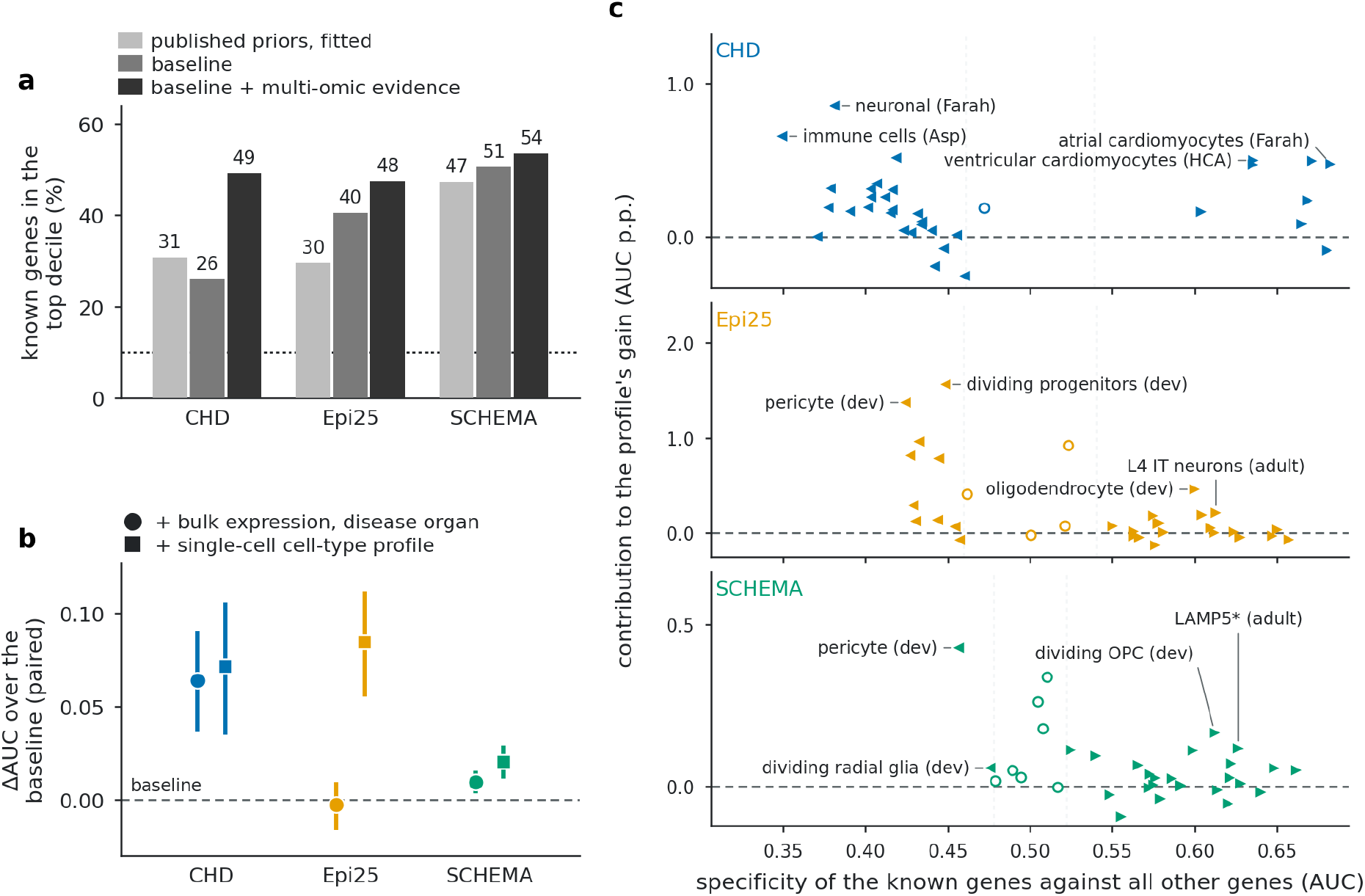
Multi-omics evidence in three case-control cohorts: gain in known genes, bulk versus single-cell expression, and contributing cell types. **(a)** Comparison of published gene priors (DOMINO, shet/GeneBayes, pHaplo and GeVIR), the baseline model and the baseline model plus multi-omics evidence (re-rank) on the three evaluated cohorts. Y-axis denotes the percentage of known disease genes included in the top decile ranking of each method. Dotted line denotes the percentage expected by chance, that is, 10% of the cohort’s curated positives. All scores are out of fold (**Additional file 1: Table S7**). **(b)** ΔAUC over the same baseline of two expression representations: the disease organ’s bulk expression level (circles) and its single-cell profile (34 heart or 33 cortex expression profiles, squares). Contribution is measured independently per cohort. **(c)** The gain in **(b)** split over cell types (Monte-Carlo Shapley, 100 random orders, shares sum to the gain). X-axis denotes the AUC with which each cell type’s specificity column separates the cohort’s known genes from all other genes and Y-axis its contribution in AUC percentage points. Left triangle: known genes less specific to that cell type than other genes, right triangle: more specific, hollow circle: AUC not distinguishable from 0.5. The two largest contributors on each side are named (LAMP5*: LAMP5 interneurons). **Additional file 1: Figs. S5-S7 and Tables S6 and S7** extend these results.

#### Expression carries most of the added multi-omics signal

To identify which feature groups accounted for the gain in AUC, we performed two complementary analyses: (a) removing each feature group from the full model and (b) adding each feature individually to the baseline model. Bulk multi-organ expression was the most informative feature group across all three cohorts: removing it reduced AUC by 0.024, 0.026 and 0.019 in CHD, Epi25 and SCHEMA, respectively. On the other hand, adding expression alone increased AUC by 0.091, 0.049 and 0.026, indicating that expression accounted for most of the incremental signal among the evaluated multi-omics features (**Additional file 1: Tables S3** and **S4**). The remaining feature groups showed negligible contribution (**Additional file 1: Fig. S4a**). For example, tissue eQTL features had sparse coverage (0.20-0.23) and performed near chance where available (AUC 0.46-0.53), whereas PTM density showed standalone discrimination (AUC 0.52-0.66) but added no signal over the baseline. The expression signal was also disease- and organ-specific (**Additional file 1: Tables S5 and S6**).

In a complementary experiment restricted to curated Mendelian disease genes (using cohort-specific disease genes as positives and genes associated with other diseases as negatives), multi-omics features still separated the two groups (**Additional file 1: Fig. S5a**), with most of the contribution coming from expression. This result is consistent with previously reported AUCs of approximately 0.7 for healthy-tissue expression as a discriminator of disease genes at tissue and cell-type resolution [41,42].

#### Single-cell expression improves discrimination

Next, we asked whether single-cell expression adds information beyond bulk expression. We compared two representations, each added independently to the baseline model: bulk expression from the disease-matched human organ and a single-cell profile comprising 34 heart or 33 cortex cell-type features across developmental stages (**Figure 4b**). Bulk expression changed AUC by +0.064 in CHD, -0.003 in Epi25 and +0.009 in SCHEMA, whereas the single-cell profile increased AUC by 0.072, 0.085 and 0.021, respectively. Single-cell expression therefore performed similarly to bulk expression in CHD and provided greater discrimination in Epi25 and SCHEMA.

Evaluating cell types individually showed that no single cell type accounted for more than 21% of the observed gain (**Figure 4c**), indicating that the information was distributed across several correlated cell types rather than dominated by one [43]. A fraction of the observed contribution came from cell types in which known disease genes showed lower specificity than other genes. The model therefore exploited both signals: reduced specificity in pericytes, immune and dividing cells, and increased specificity in cardiomyocytes for CHD and in neurons for Epi25 and SCHEMA. This pattern is consistent with many Mendelian disease genes being broadly expressed rather than restricted to the affected tissue, such that no single cell type alone separates disease genes from other genes [41,44].

#### Concordance with the cohort’s own burden signal and model transfer across cohorts

We next asked whether genes highly ranked by a model that included multi-omics features were also more likely to show significant burden associations in each cohort. The model was trained on curated disease-gene labels, excluding burden statistics and all other cohort-derived features. We then tested whether genes in the top 10% of the resulting ranking were enriched for nominally significant burden associations (P < 0.05). We observed significant enrichment in all three cohorts: CHD (*P* = 1.9 × 10⁻⁴, *OR* = 1.41), Epi25 (*P* = 5.3 × 10⁻⁴, *OR* = 1.28) and SCHEMA (*P* = 6.2 × 10⁻³⁸, *OR* = 2.56) (**Figure 5a-c**). These analyses were adjusted for gene mutational target size. Although burden statistics were excluded from the model, its curated training labels may reflect prior association evidence, so the two sources are not fully independent. We also tested whether a model fitted on one cohort could be applied to another. In every valid comparison, the transferred model underperformed the best published gene-prior score in the target cohort (ΔAUC = -0.162 to -0.103) (**Figure 5d**).

**Figure 5.**
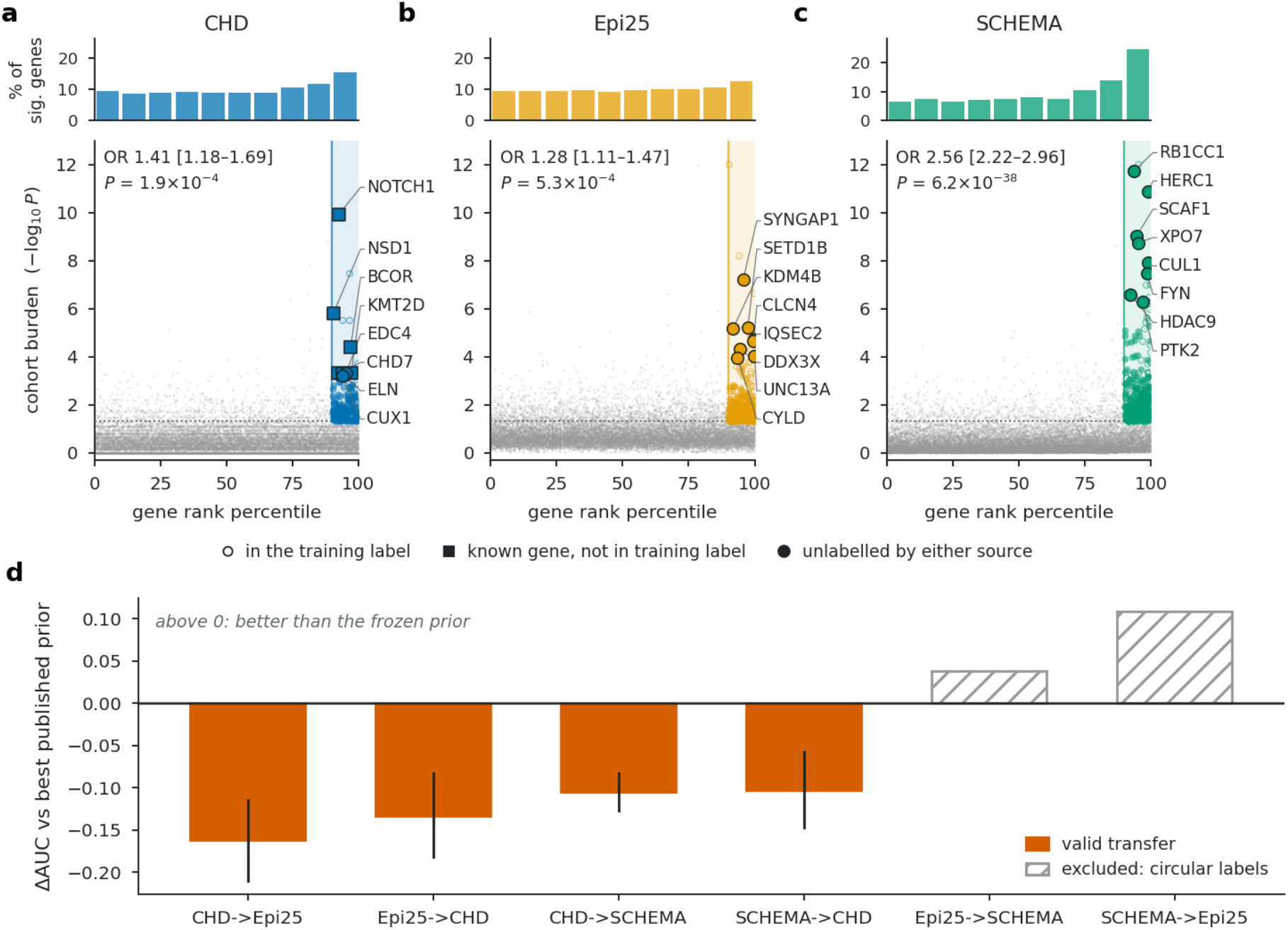
Concordance with the burden signal of the own cohort, and cross-cohort transfer without refitting. (a-c) Each grey point is one gene. X-axis denotes the gene’s percentile in a cohort-blind ranking, fitted out of fold on constraint and multi-omics features with no access to the burden statistic. Y-axis denotes the cohort’s burden evidence and the dotted line the nominally significant p-value = 0.05. Shaded band represents the top decile of the cohort-blind ranking. The histogram above each panel gives the share of nominally burden-significant genes falling in each decile of that ranking. Odds ratio, 95% confidence interval and p-value give the enrichment of genes reaching burden P<0.05 in the top decile, adjusted for mutational target size. Selected genes in the top ranking decile with strong burden associations are labelled for illustration. **(d)** ΔAUC of the same model fitted on one cohort applied unchanged to another, measured against the best published prior on the target cohort. Hatched pairs are excluded rather than measured: the SCHEMA label as published contains the Epi25 label (**Additional file 1: Fig. S4b**).

Taken together, adding multi-omic evidence assembled from different resources improves ranking beyond what the published priors achieve, and the gain is disease-specific and organ-matched. Applied to cohort-derived burden statistics, hvantk re-ranking produces a per-gene score that improves interpretation rather than the discovery of novel candidate genes.

## Discussion

hvantk addresses the three challenges outlined above. Provenance is handled by recording dataset versions and detecting source-specific changes, making upstream updates traceable as reference databases and datasets evolve [36]. Extensibility is provided through a validated plugin system, allowing new data sources, disease cohorts and feature sets to be added and analysed without changing the framework core code. Dedicated plugin and workflow skills enable researchers and AI coding agents to develop data plugins and execute analysis workflows, while the underlying computations remain deterministic and independent of large language models (LLMs). Finally, a common data model extending Hail *MatrixTable* and AnnData allows cohort-scale stages to run on Spark or HPC cluster and downstream variant- and gene-level analyses to run locally when the data fit in memory.

hvantk differs from emerging agentic genomics frameworks [16] because the AI agent authors data plugins and drives analysis workflows through skills, whereas the analysis itself is executed by deterministic tools. It complements modular annotation tools [37] by linking variant-, gene- and protein-level evidence to cohort-level association results. It also differs from gene-prior scores [38,39], which rank genes without taking into account the allelic architecture and burden statistics observed in a given cohort.

We evaluated five open- and closed-source AI coding models across three data-plugin development tasks (HGNC, UCSC Cell Browser and dbNSFP) to assess their ability to generate plugins that pass the unit tests (**Figure 3c-e**). Success rates, token consumption and execution time varied across models and tasks. In this benchmark, Claude models generally required fewer iterations to pass the tests, highlighting the importance of model selection for reliably and efficiently extending the annotation framework with new data sources.

The workflow measures what multi-omics annotation adds to burden interpretation rather than to statistical discovery. The burden statistic and its significance remain unchanged, whereas the added information is used to re-rank genes across the tested gene universe. In the evaluated cohorts, bulk and single-cell expression added signal over a baseline that included burden statistics, gene constraint and publication count. This suggests that additional omics features should be evaluated for incremental value relative to a well-defined baseline. PTM density illustrates this point: it showed a modest AUC of 0.66, but provided no additional predictive value beyond gene constraint. In contrast, the eQTL result should be interpreted with caution due to its sparse coverage (0.20-0.23 of genes). The eQTL catalogue used in this study derives from adult GTEx tissues, and eQTL evidence for the developmental and disease-specific contexts of these cohorts remains limited. The observed weak signal may therefore reflect the current state of the resource rather than the value of eQTL evidence as such.

The added expression signal was disease-specific and strongest in the corresponding organ across all three cohorts. Among curated Mendelian disease genes, AUCs of 0.71-0.81 were of the same order as values reported for healthy-tissue expression as a discriminator of disease genes [41,42]. However, the studies differ in labels, gene universes and evaluation design. The distributed single-cell contribution across cell types is also consistent with previous mapping of schizophrenia genes to several neuronal classes rather than a single one [43]. Because the positive and the negative genes were derived from ClinGen and GenCC, curation bias remains a possible explanation. However, the most informative organ changed with disease - heart for CHD and brain for Epi25 and SCHEMA - arguing against a single tissue bias shared across cohorts. Independent CHD studies based on *de novo* variant enrichment and cardiac tissue profiling support the same biological pattern [45,46].

This study has several limitations. Cohorts matching the same disease with sufficient power for replication were not available, and CHD estimates showed broader confidence intervals given 234 curated positives among 16,349 genes. The positive set is incomplete and correlated with both constraint and publication count, so that undiscovered disease genes may be labelled as negatives. Finally, the allelic-architecture and quality-control assessment requires variant-level cohort data. Cohorts providing gene-level summary data only cannot apply it directly, and a single problematic recurrent variant may still distort the signal.

## Conclusions

hvantk provides an interoperable framework linking cohort-scale joint genotyping, burden analysis, provenance-aware multi-omics annotation and gene re-ranking. By extending Hail and AnnData data structures, its common data representations support distributed and local analyses. In addition, schema-validated data plugins and dedicated workflow skills enable AI coding agents to integrate new resources and orchestrate analyses, while dataset generation and analytical computations remain independent of large language models (LLMs).

The analysis of three case-control cohorts (CHD, Epi25 and SCHEMA) showed that adding multi-omics features to burden statistics, gene constraint and publication count improved the ranking of curated disease genes relative to this baseline and published gene-prior scores. Bulk and single-cell expression contributed most of the additional signal, with patterns varying by disease and organ, whereas eQTL and PTM features contributed to a lesser degree. hvantk provides a framework for evaluating the contribution of new annotation resources as they become available. We expect this iterative exchange between public multi-omics resources and in-house cohorts, rather than any single annotation source, to drive further gains in rare-variant gene interpretation.

## Methods

### Framework and data provenance

hvantk represents datasets using four defined types: AnnotationTable, ExpressionMatrix, VariantMatrix and GeneSet. Each dataset carries a provenance record containing the source plugin, dataset identifier, plugin version, upstream source fingerprint, schema identifier, build timestamp and provenance of parent datasets. Shared biological identifiers support joins across annotation levels while retaining the provenance of contributing datasets. Depending on data structure and scale, hvantk uses Hail Tables and MatrixTables for distributed variant and cohort data [2], AnnData for expression matrices and pandas for in-memory tabular analyses. The package structure is described in the *hvantk* documentation (https://github.com/bigbio/hvantk).

### Data sources and source monitoring

Twenty-one public resources (among variant annotation, gene-level information, molecular QTLs, protein features and expression data sources) were integrated into hvantk (**Table 1**), generating a total of 26 datasets. Each provider is implemented as a data plugin that retrieves or imports source data, processes and converts them into the hvantk data model. Fifteen providers include automated downloaders, whereas acquisition procedures are documented for the remaining ones.

Datasets implemented in hvantk record the source version and an upstream fingerprint at build time, used later to detect source changes. All 26 datasets include a reference schema and representative row snapshot for regression testing, and a live upstream check against a previous successful build. A scheduled workflow performs these checks at configurable intervals and opens draft pull requests when changes are detected. Synthetic test data are used when source licensing restrictions do not allow redistribution of original data (**Additional file 1: Fig. S3**).

### AI-assisted plugin development benchmark

Five AI coding models (Claude Sonnet 5.5, Claude Haiku 4.5, DeepSeek-V4-Flash, Qwen3-Coder-Next and GLM-4.7-Flash) were evaluated on three plugin development tasks representing different data sources (HGNC, UCSC Cell Browser and dbNSFP). Each model/provider combination was evaluated in three independent runs, allowing up to 10 iterations per run. The coding agent implemented a data plugin per iteration and validated it against the plugin specification and automated tests, using test failures as feedback. Success was defined as passing the required unit test within the iteration limit. The number of iterations required for success, token consumption and wall-clock execution time were summarized across the three runs for each model/provider combination.

### Joint genotyping and scalability benchmarking

#### Cohort assembly

Joint genotyping was performed with the *hvantk hgc* pipeline using resumable processing stages. Single-sample gVCFs were combined into a Hail Variant Dataset (VDS) [2] and densified for downstream analysis, multi-allelic sites were split, and a multi-sample VCF was exported with allele counts and frequencies recalculated from the joint callset. Quality control included per-sample call rate, transition/transversion ratio, mean depth and genotype quality, and per-variant allele counts, allele frequencies, call rate and Hardy-Weinberg equilibrium statistics.

#### Scalability benchmarking

Performance was evaluated using whole-genome gVCFs from 1,005 samples of the primary CHD cohort [10] on a Spark cluster. Unless otherwise stated, benchmarks were restricted to chromosome 1. We tested computational scaling at a fixed cohort size (n=500, 16-128 cores), cohort-size scaling at 48 cores (n=50-1,005), and an additional chromosome 20 experiment to separate effects of cohort size from partition number.

### Disease cohorts, gene universes and training labels

#### Disease cohorts

Three case-control cohorts were analysed. The primary congenital heart disease (CHD) exome cohort [10] was processed using the *hvantk hgc* workflow. A subset of 1,005 samples resequenced by whole-genome sequencing was used exclusively for scalability benchmarking, whereas all downstream CHD analyses used the exome dataset. Two additional cohorts were included: the Epi25 developmental and epileptic encephalopathy cohort [8] and the SCHEMA schizophrenia meta-analysis [9].

The cohorts differed in the level of data available for downstream analysis. CHD was represented by de-identified variant-level summaries containing consequence classes, pathogenicity scores, population allele frequencies and carrier counts by cohort stratum, together with per-gene burden statistics. SCHEMA provided published gene-level burden statistics and per-variant case/control allele counts, allowing assessment of allelic architecture. Epi25 was available only at the gene level and was therefore excluded from the allelic-architecture assessment. Gene re-ranking was performed using per-gene burden statistics from all three cohorts, without requiring individual-level genotypes.

#### Gene universes and disease labels

The primary evaluation included all genes with available burden statistics, without filtering on burden significance: 16,349 genes for CHD (**Additional file 2**), 15,956 for Epi25 (**Additional file 3**) and 18,569 for SCHEMA (**Additional file 4**). Positive labels were assigned to genes with Definitive, Strong or Moderate evidence for the corresponding disease in ClinGen [24] or GenCC [25], accessed on 23 March 2026. This resulted in 234, 200 and 710 positive genes for CHD, Epi25 and SCHEMA, respectively. Genes without qualifying curated associations were treated as negatives for model training and evaluation. However, these constitute an unlabelled set that may include undiscovered disease genes. Models were therefore evaluated against the observed curated disease-gene labels rather than independently verified negative controls. A broader literature-derived CHD gene panel was additionally used for sensitivity analysis.

Disease-specific discrimination was evaluated separately within the subset of curated Mendelian disease genes, comprising 3,952, 3,871 and 4,157 genes for CHD, Epi25 and SCHEMA, respectively. In this analysis, genes associated with the corresponding cohort disease were treated as positives and genes associated with other diseases as negatives.

### Multi-omics features

Gene-level features were assembled from a common catalogue comprising gene constraint metrics (gnomAD LOEUF and MOEUF [22]), expression, eQTL information, post-translational modification (PTM) density, allelic architecture, burden statistics and variant-to-gene prediction score aggregates. Expression features included developmental bulk transcriptomes from six organs (brain, heart, kidney, liver, ovary and testis) in humans and mice [47], GTEx tissue-level expression [27] and developmental single-cell atlases [48,49]. Disease-matched single-cell profiles comprised 34 heart cell-type features for CHD and 33 cortex cell-type features for Epi25 and SCHEMA. PTM density was derived by mapping UniProt modification sites to genomic codon positions [30]. Allelic-architecture and quality-control features were excluded from the supervised re-ranking. Protein-interaction network proximity was not considered to avoid potential circularity from curated disease-gene labels.

### Gene re-ranking and model validation

A gradient-boosting classifier was used to implement the model for gene re-ranking. The baseline model included cohort-specific burden statistics, gnomAD constraint metrics (LOEUF and MOEUF) and publication count. In addition, models with multi-omics evidence considered expression, eQTL and PTM features. Models were evaluated using fivefold group cross-validation, grouping genes by their primary HGNC gene group to reduce information leakage. Across 40 alternative fold assignments, multi-omics features improved AUC over the baseline in every cohort (ΔAUC +0.061 to +0.113 in CHD, +0.039 to +0.087 in Epi25 and +0.024 to +0.035 in SCHEMA). All reported predictions were generated out of fold.

To compare performance, four published gene-prior scores (GeVIR [23], DOMINO [38], shet [39] and pHaplo [40]) were evaluated for their ability to distinguish curated disease-associated genes from other genes in each cohort. Scores were assessed individually and combined by fitting a gradient-boosting classifier, using the same gene universes, labels and cross-validation partitions as the baseline and multi-omics models. GeVIR percentiles were inverted, and missing prior values were imputed using the cohort median.

### Evaluation and complementary analysis

Accuracy on the classification was assessed using out-of-fold ROC-AUC and average precision (AP), with paired 95% confidence intervals calculated from 2,000 gene-level bootstrap resamples. Feature contributions were evaluated by removing each group from the full model or adding it individually to the baseline model. Single-cell and bulk-level expression contributions were measured independently against the same baseline, and the single-cell gain was split over cell types by Monte-Carlo Shapley attribution (100 random orders).

A separate model trained without burden statistics or other cohort-derived features was used to evaluate concordance between the re-ranking results and the cohort’s own burden evidence. Enrichment of nominally burden-significant genes (P < 0.05) in the top ranked decile was assessed using odds ratios adjusted for mutational target size. Cross-disease transfer was evaluated by applying models trained on gene-intrinsic features to a second cohort without refitting. Cohort pairs with positive-label overlap of at least 50% of the smaller positive set, such as the Epi25/SCHEMA pair, were excluded.

## Declarations

### Ethics approval and consent to participate

The primary CHD cohort’s ethics approval and participant consent are covered by the companion study [10]. Access to congenital heart disease (CHD) case data was granted by the European Genome-phenome Archive (EGA) Data Access Committees following protocol review. The datasets used in the study had prior institutional research ethics board approval from the German Heart Registry (Berlin, Germany), the University of Leuven, Belgium (number: B322201010111/S52853), and the University of Nottingham by the East Midlands-Leicester Research Ethics Committee (number 6721). The DDD study was approved by the UK Research Ethics Committee (10/H0305/83, granted by the Cambridge South Research Ethics Committee and GEN/284/12, granted by the Republic of Ireland Research Ethics Committee). UK Biobank whole-exome sequencing data were accessed under approved application number 44165.

The Epi25 and SCHEMA analyses were based strictly on previously published, publicly available variant- and gene-level summary statistics released by the original study consortia [8,9]. No individual-level sequencing, genotype or clinical data from these cohorts were accessed. These analyses therefore represent secondary use of publicly released, aggregated genetic association data.

### Consent for publication

Not applicable.

### Data and code availability

The public data sources integrated by hvantk are listed in **Table 1** and are available from their respective providers. Burden statistics for the three cohorts analysed in the present study are available from their original publications. All further data supporting this work are included within the article and its additional files. Code availability: https://github.com/bigbio/hvantk.

### Authors contributions

E.A. and Y.P.-R. designed and implemented the framework, analyses and drafted the initial manuscript. Y.P.-R. supervised the software architecture and data-integration design. M.-P.H. supervised the study. All authors read and approved the final manuscript.

## Acknowledgements

The benchmarks and cohort analyses were conducted on the HPC cluster ROSA, located at the University of Oldenburg (Germany). ROSA was funded by the Deutsche Forschungsgemeinschaft (DFG, German Research Foundation) through its Major Research Instrumentation Programme - Project-ID 493597399 (INST 184/225-1 FUGG) and the Ministry of Science and Culture (MWK) of Lower Saxony.

## Supplementary Information

**Additional file 1:** Figures S1-S7 and Tables S1-S7. Supplementary figures and tables with legends.

**Additional file 2:** Per-gene (n=16,349) re-ranking scores for the congenital heart disease (CHD) cohort with the out-of-fold scores, curated disease-gene labels, burden statistic and published gene-prior scores. Columns are described in the legend sheet.

**Additional file 3:** Per-gene (n=15,956) re-ranking scores for the developmental and epileptic encephalopathy (Epi25) cohort with the same columns as Additional file 2.

**Additional file 4:** Per-gene (n=18,569) re-ranking scores for the schizophrenia (SCHEMA) cohort with the same columns as Additional file 2.

## References

[1] Hanssen F, Garcia MU, Folkersen L, Pedersen AS, Lescai F, Jodoin S, et al. Scalable and efficient DNA sequencing analysis on different compute infrastructures aiding variant discovery. NAR Genomics and Bioinformatics 2024;6(2):lqae031. doi:10.1093/nargab/lqae031.

[2] Poterba T, Vittal C, King D, Goldstein D, Goldstein JI, Schultz P, et al. The scalable variant call representation: Enabling genetic analysis beyond one million genomes. Bioinformatics 2025;41(1):btae746. doi:10.1093/bioinformatics/btae746.

[3] Chen S, Francioli LC, Goodrich JK, Collins RL, Kanai M, Wang Q, et al. A genomic mutational constraint map using variation in 76,156 human genomes. Nature 2024;625(7993):92–100. doi:10.1038/s41586-023-06045-0.

[4] Perez-Riverol Y, Bittremieux W, Noble WS, Martens L, Bilbao A, Lazear MR, et al. Open-source and FAIR research software for proteomics. Journal of Proteome Research 2025;24(5):2222–2234. doi:10.1021/acs.jproteome.4c01079.

[5] Xu M, Yan J, Feng R, Cai Q, Zhang P, Zhao C, et al. BioClaw: Human-bot research collaboration ecosystems in group chats. bioRxiv 2026. doi:10.64898/2026.04.11.716807.

[6] Poplin R, Ruano-Rubio V, DePristo MA, Fennell TJ, Carneiro MO, Van der Auwera GA, et al. Scaling accurate genetic variant discovery to tens of thousands of samples. bioRxiv 2017. doi:10.1101/201178.

[7] Virshup I, Rybakov S, Theis FJ, Angerer P, Wolf FA. anndata: Access and store annotated data matrices. Journal of Open Source Software 2024;9(101):4371. doi:10.21105/joss.04371.

[8] Epi25 Collaborative. Exome sequencing of 20,979 individuals with epilepsy reveals shared and distinct ultra-rare genetic risk across disorder subtypes. Nature Neuroscience 2024;27(10):1864–1879. doi:10.1038/s41593-024-01747-8.

[9] Singh T, Poterba T, Curtis D, Akil H, Al Eissa M, Barchas JD, et al. Rare coding variants in ten genes confer substantial risk for schizophrenia. Nature 2022;604(7906):509–516. doi:10.1038/s41586-022-04556-w.

[10] Audain E, Wilsdon A, Dombrowsky G, Sifrim A, Breckpot J, Perez-Riverol Y, et al. Assessing the contribution of rare variants to congenital heart disease through a large-scale case-control exome study. npj Genomic Medicine 2026;11(1):30. doi:10.1038/s41525-026-00582-z.

[11] Mbatchou J, Barnard L, Backman J, Marcketta A, Kosmicki JA, Ziyatdinov A, et al. Computationally efficient whole-genome regression for quantitative and binary traits. Nature Genetics 2021;53(7):1097–1103. doi:10.1038/s41588-021-00870-7.

[12] Li X, Li Z, Zhou H, Gaynor SM, Liu Y, Chen H, et al. Dynamic incorporation of multiple in silico functional annotations empowers rare variant association analysis of large whole-genome sequencing studies at scale. Nature Genetics 2020;52(9):969–983. doi:10.1038/s41588-020-0676-4.

[13] Balachandran S, Prada-Medina CA, Mensah MA, Kakar N, Nagel I, Pozojevic J, et al. STIGMA: Single-cell tissue-specific gene prioritization using machine learning. The American Journal of Human Genetics 2024;111(2):338–349. doi:10.1016/j.ajhg.2023.12.011.

[14] Madrigal P, Thanki AS, Fexova S, Yu ID, Chatzigeorgiou A, Zucchi I, et al. Expression Atlas in 2026: Enabling FAIR and open expression data through community collaboration and integration. Nucleic Acids Research 2026;54(D1):D147–D157. doi:10.1093/nar/gkaf1238.

[15] Desiere F, Deutsch EW, King NL, Nesvizhskii AI, Mallick P, Eng J, et al. The PeptideAtlas project. Nucleic Acids Research 2006;34(suppl_1):D655–D658. doi:10.1093/nar/gkj040.

[16] Corpas M, Guio H, Fatumo S. Agentic genomics: From pipeline automation to autonomous validation. Cell Genomics 2026;6(8):101305. doi:10.1016/j.xgen.2026.101305.

17. 1000 Genomes Project Consortium. A global reference for human genetic variation. Nature 2015;526(7571):68–74. doi:10.1038/nature15393.

[18] Landrum MJ, Chitipiralla S, Kaur K, Brown G, Chen C, Hart J, et al. ClinVar: Updates to support classifications of both germline and somatic variants. Nucleic Acids Research 2025;53(D1):D1313–D1321. doi:10.1093/nar/gkae1090.

[19] Liu X, Li C, Mou C, Dong Y, Tu Y. dbNSFP v4: A comprehensive database of transcript-specific functional predictions and annotations for human nonsynonymous and splice-site SNVs. Genome Medicine 2020;12(1):103. doi:10.1186/s13073-020-00803-9.

[20] Avsec Ž, Latysheva N, Cheng J, Novati G, Taylor KR, Ward T, et al. Advancing regulatory variant effect prediction with AlphaGenome. Nature 2026;649(8099):1206–1218. doi:10.1038/s41586-025-10014-0.

[21] Cerezo M, Sollis E, Ji Y, Lewis E, Abid A, Bircan KO, et al. The NHGRI-EBI GWAS Catalog: Standards for reusability, sustainability and diversity. Nucleic Acids Research 2025;53(D1):D998–D1005. doi:10.1093/nar/gkae1070.

[22] Karczewski KJ, Francioli LC, Tiao G, Cummings BB, Alföldi J, Wang Q, et al. The mutational constraint spectrum quantified from variation in 141,456 humans. Nature 2020;581(7809):434–443. doi:10.1038/s41586-020-2308-7.

[23] Abramovs N, Brass A, Tassabehji M. GeVIR is a continuous gene-level metric that uses variant distribution patterns to prioritize disease candidate genes. Nature Genetics 2020;52(1):35–39. doi:10.1038/s41588-019-0560-2.

[24] Rehm HL, Berg JS, Brooks LD, Bustamante CD, Evans JP, Landrum MJ, et al. ClinGen: The Clinical Genome Resource. New England Journal of Medicine 2015;372(23):2235–2242. doi:10.1056/nejmsr1406261.

[25] DiStefano MT, Goehringer S, Babb L, Alkuraya FS, Amberger J, Amin M, et al. A global effort to harmonize gene-disease evidence resources. Genetics in Medicine 2022;24(8):1732–1742. doi:10.1016/j.gim.2022.04.017.

[26] Sondka Z, Dhir NB, Carvalho-Silva D, Jupe S, Madhumita, McLaren K, et al. COSMIC: A curated database of somatic variants and clinical data for cancer. Nucleic Acids Research 2024;52(D1):D1210–D1217. doi:10.1093/nar/gkad986.

[27] GTEx Consortium. The GTEx Consortium atlas of genetic regulatory effects across human tissues. Science 2020;369(6509):1318–1330. doi:10.1126/science.aaz1776.

[28] Fang H, Jiang L, da Veiga Leprevost F, Jian R, Chan J, Glinos D, et al. Regulation of protein abundance in normal human tissues. medRxiv 2025. doi:10.1101/2025.01.10.25320181.

[29] Meyer MJ, Beltrán JF, Liang S, Fragoza R, Rumack A, Liang J, et al. Interactome INSIDER: A structural interactome browser for genomic studies. Nature Methods 2018;15(2):107–114. doi:10.1038/nmeth.4540.

[30] UniProt Consortium. UniProt: The Universal Protein Knowledgebase in 2025. Nucleic Acids Research 2025;53(D1):D609–D617. doi:10.1093/nar/gkae1010.

[31] Speir ML, Bhaduri A, Markov NS, Moreno P, Nowakowski TJ, Papatheodorou I, et al. UCSC Cell Browser: Visualize your single-cell data. Bioinformatics 2021;37(23):4578–4580. doi:10.1093/bioinformatics/btab503.

[32] Edwards NJ, Oberti M, Thangudu RR, Cai S, McGarvey PB, Jacob S, et al. The CPTAC data portal: A resource for cancer proteomics research. Journal of Proteome Research 2015;14(6):2707–2713. doi:10.1021/pr501254j.

[33] Liberzon A, Birger C, Thorvaldsdóttir H, Ghandi M, Mesirov JP, Tamayo P. The Molecular Signatures Database (MSigDB) hallmark gene set collection. Cell Systems 2015;1(6):417–425. doi:10.1016/j.cels.2015.12.004.

[34] Yates AD, Austine-Orimoloye O, Azov AG, Barba M, Barnes I, Barrera-Enriquez VP, et al. Ensembl 2026. Nucleic Acids Research 2026;54(D1):D1053–D1060. doi:10.1093/nar/gkaf1239.

[35] Seal RL, Braschi B, Gray K, McClay J, Tweedie S, Bruford EA. Genenames.org: The HGNC and PGNC resources in 2026. Nucleic Acids Research 2026;54(D1):D1098–D1107. doi:10.1093/nar/gkaf1229.

[36] Li L, Tian X, Woodzell V, Gibbs RA, Yuan B, Venner E. Tracking updates in clinical databases increases efficiency for variant reanalysis. Genetics in Medicine Open 2024;2:101841. doi:10.1016/j.gimo.2024.101841.

[37] McLaren W, Gil L, Hunt SE, Riat HS, Ritchie GR, Thormann A, et al. The Ensembl Variant Effect Predictor. Genome Biology 2016;17(1):122. doi:10.1186/s13059-016-0974-4.

[38] Quinodoz M, Royer-Bertrand B, Cisarova K, Di Gioia SA, Superti-Furga A, Rivolta C. DOMINO: Using machine learning to predict genes associated with dominant disorders. The American Journal of Human Genetics 2017;101(4):623–629. doi:10.1016/j.ajhg.2017.09.001.

[39] Zeng T, Spence JP, Mostafavi H, Pritchard JK. Bayesian estimation of gene constraint from an evolutionary model with gene features. Nature Genetics 2024;56(8):1632–1643. doi:10.1038/s41588-024-01820-9.

[40] Collins RL, Glessner JT, Porcu E, Lepamets M, Brandon R, Lauricella C, et al. A cross-disorder dosage sensitivity map of the human genome. Cell 2022;185(16):3041–3055.e25. doi:10.1016/j.cell.2022.06.036.

[41] Feiglin A, Allen BK, Kohane IS, Kong SW. Comprehensive analysis of tissue-wide gene expression and phenotype data reveals tissues affected in rare genetic disorders. Cell Systems 2017;5(2):140–148.e2. doi:10.1016/j.cels.2017.06.016.

[42] Novoa J, Pazos F, Chagoyen M. Exploring cell-type associations of rare disease phenotypes using non-diseased cell atlases. Briefings in Bioinformatics 2026;27(4):bbag426. doi:10.1093/bib/bbag426.

[43] Skene NG, Bryois J, Bakken TE, Breen G, Crowley JJ, Gaspar HA, et al. Genetic identification of brain cell types underlying schizophrenia. Nature Genetics 2018;50(6):825–833. doi:10.1038/s41588-018-0129-5.

[44] Hekselman I, Yeger-Lotem E. Mechanisms of tissue and cell-type specificity in heritable traits and diseases. Nature Reviews Genetics 2020;21(3):137–150. doi:10.1038/s41576-019-0200-9.

[45] Sierant MC, Jin SC, Bilguvar K, Morton SU, Dong W, Jiang W, et al. Genomic analysis of 11,555 probands identifies 60 dominant congenital heart disease genes. Proceedings of the National Academy of Sciences 2025;122(13):e2420343122. doi:10.1073/pnas.2420343122.

[46] Hill MC, Kadow ZA, Long H, Morikawa Y, Martin TJ, Birks EJ, et al. Integrated multi-omic characterization of congenital heart disease. Nature 2022;608(7921):181–191. doi:10.1038/s41586-022-04989-3.

[47] Cardoso-Moreira M, Halbert J, Valloton D, Velten B, Chen C, Shao Y, et al. Gene expression across mammalian organ development. Nature 2019;571(7766):505–509. doi:10.1038/s41586-019-1338-5.

[48] Asp M, Giacomello S, Larsson L, Wu C, Fürth D, Qian X, et al. A spatiotemporal organ-wide gene expression and cell atlas of the developing human heart. Cell 2019;179(7):1647–1660. doi:10.1016/j.cell.2019.11.025.

[49] Farah EN, Hu RK, Kern C, Zhang Q, Lu TY, Ma Q, et al. Spatially organized cellular communities form the developing human heart. Nature 2024;627(8005):854–864. doi:10.1038/s41586-024-07171-z.

