## Additional file 1 for "hvantk: an agentic multi-omics framework for variant annotation and analysis"

**Additional file 1. Supplementary figures and tables**

**Contents**

Supplementary Figure S1. Scalability benchmark of the HGC pipeline.

Supplementary Figure S2. Quality control of the HGC joint callset.

Supplementary Figure S3. Plugin contract status of the 26 datasets.

Supplementary Figure S4. Diagnostics of the feature groups and overlap between disease labels.

Supplementary Figure S5. Cross-disease specificity and cell-type expression.

Supplementary Figure S6. Cell-type contributions to the single-cell expression gain.

Supplementary Figure S7. Bulk and single-cell expression compared at matched breadth.

Supplementary Table S1. ΔAUC of multi-omics features over three reference models.

Supplementary Table S2. Discrimination of the gene-prior scores and ranking models.

Supplementary Table S3. Contribution of each feature group by removal from the full model.

Supplementary Table S4. Coverage and discrimination of each feature group.

Supplementary Table S5. Cross-disease specificity of each feature group.

Supplementary Table S6. Discrimination of disease genes by organ expression.

Supplementary Table S7. Recovery of known disease genes by each ranking method.

Supplementary References

### Supplementary Figures

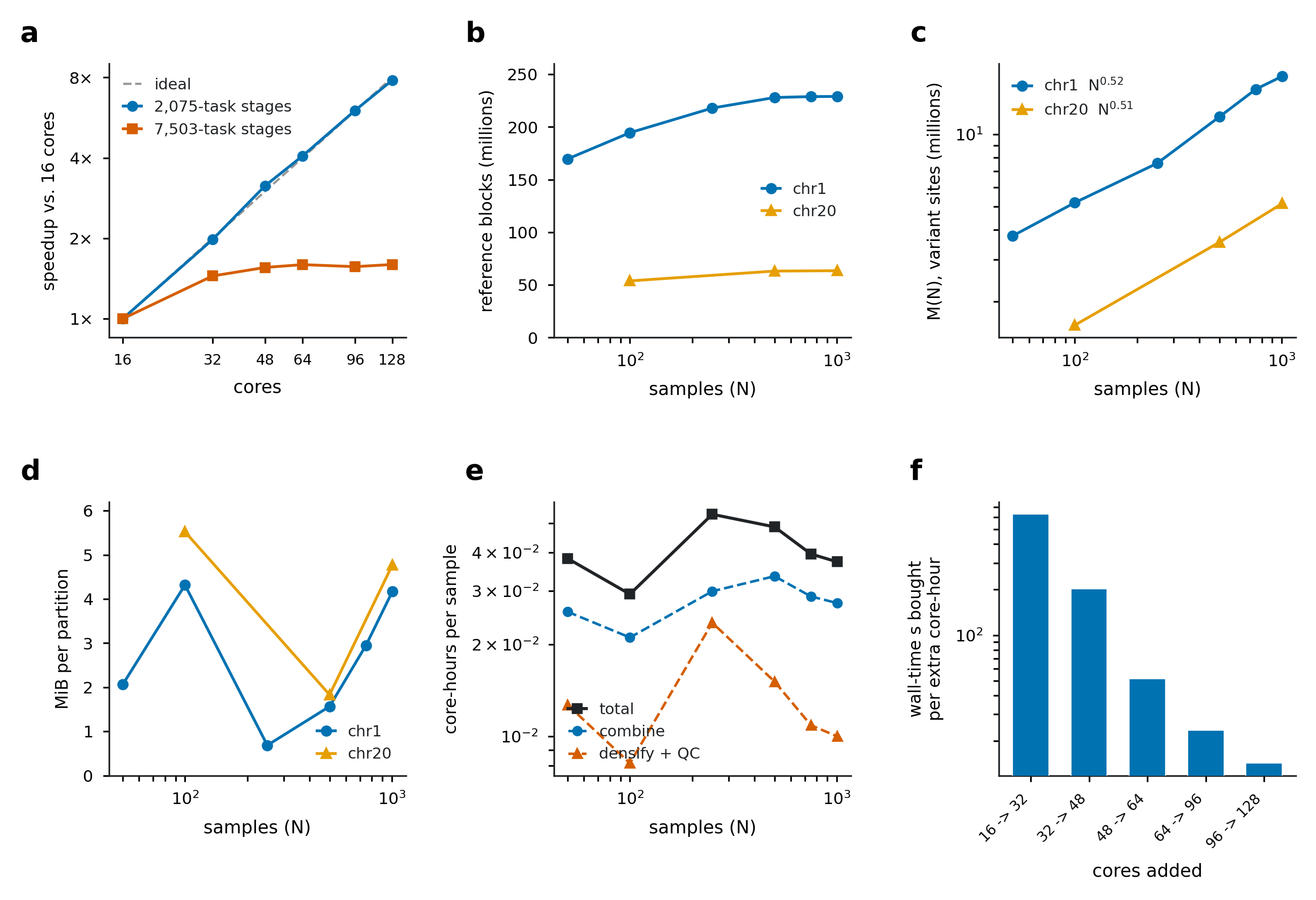

**Supplementary Figure S1. Scalability benchmark of the HGC pipeline.** Whole-genome gVCFs from a fraction (N=1,005) of the primary CHD cohort were processed on a Spark cluster, restricted to chromosome 1 unless otherwise stated. Each point represents a single run. **(a)** Speedup over 16 cores at N=500 for Spark stages grouped by number of tasks. Dashed line denotes ideal scaling. **(b)** Number of reference blocks against the number of samples (N) for chr1 and chr20. **(c)** Number of variant sites, M(N), against N, with the fitted power-law exponents. **(d)** Data per partition of the dense MatrixTable against N. **(e)** Core-hours per sample against N for the combine step, densification and quality control, and in total. **(f)** Wall-clock time saved per additional core-hour at N=500, as cores are added from 16 to 128.

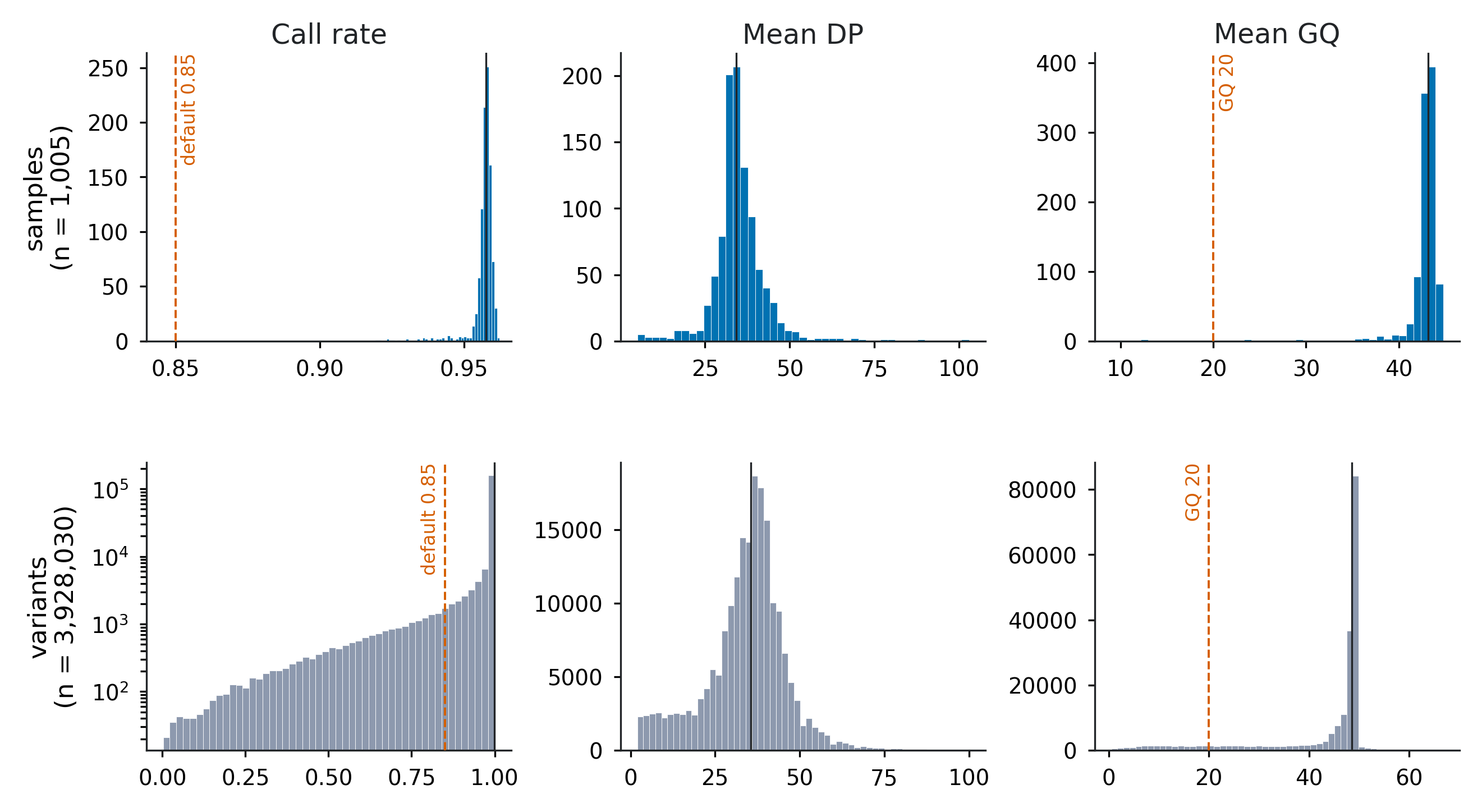

**Supplementary Figure S2. Computation and visulization of quality control (QC) metrics from the HGC joint callset.** Call rate, mean depth (DP) and mean genotype quality (GQ) computed per sample (top row, n=1,005) and per variant (bottom row, n=3,928,030 polymorphic variants) for chromosome 1. Dashed lines denote the hvantk default filter thresholds and solid lines denote the medians.

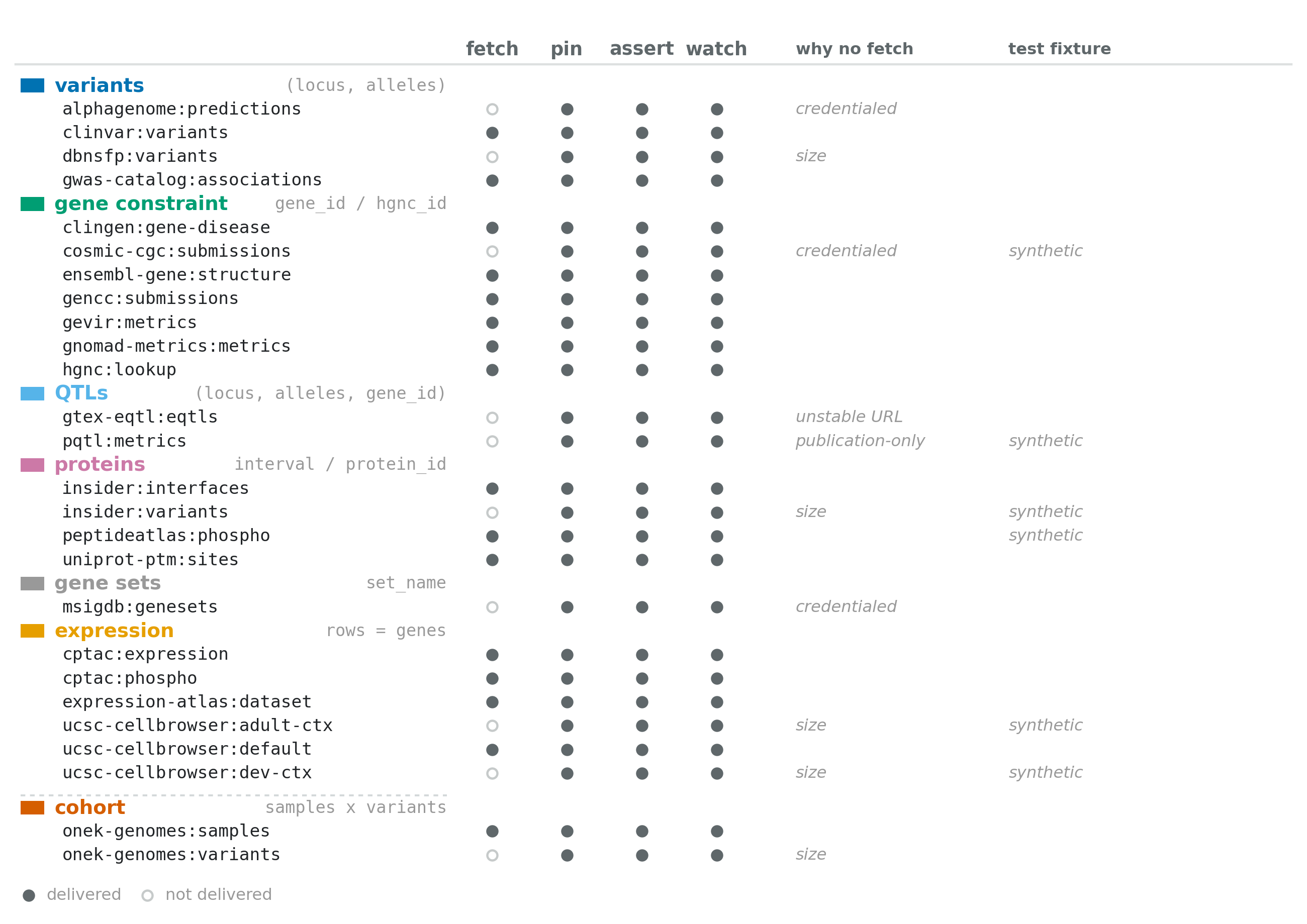

**Supplementary Figure S3. Datasets generated by the 21 data plugins of hvantk v0.6.0**. Datasets are grouped by biological data type and primary identifiers. Filled circles denote a built-in downloader (fetch, 16 datasets), a reference schema and row snapshot (pin, 26), a test comparing a fixture build against that snapshot (assert, 26) and a live upstream check (watch, 26). For datasets without a downloader, the reason why acquisition is documented instead is given. Synthetic test data are used where the source licence does not allow redistribution (test fixture column).

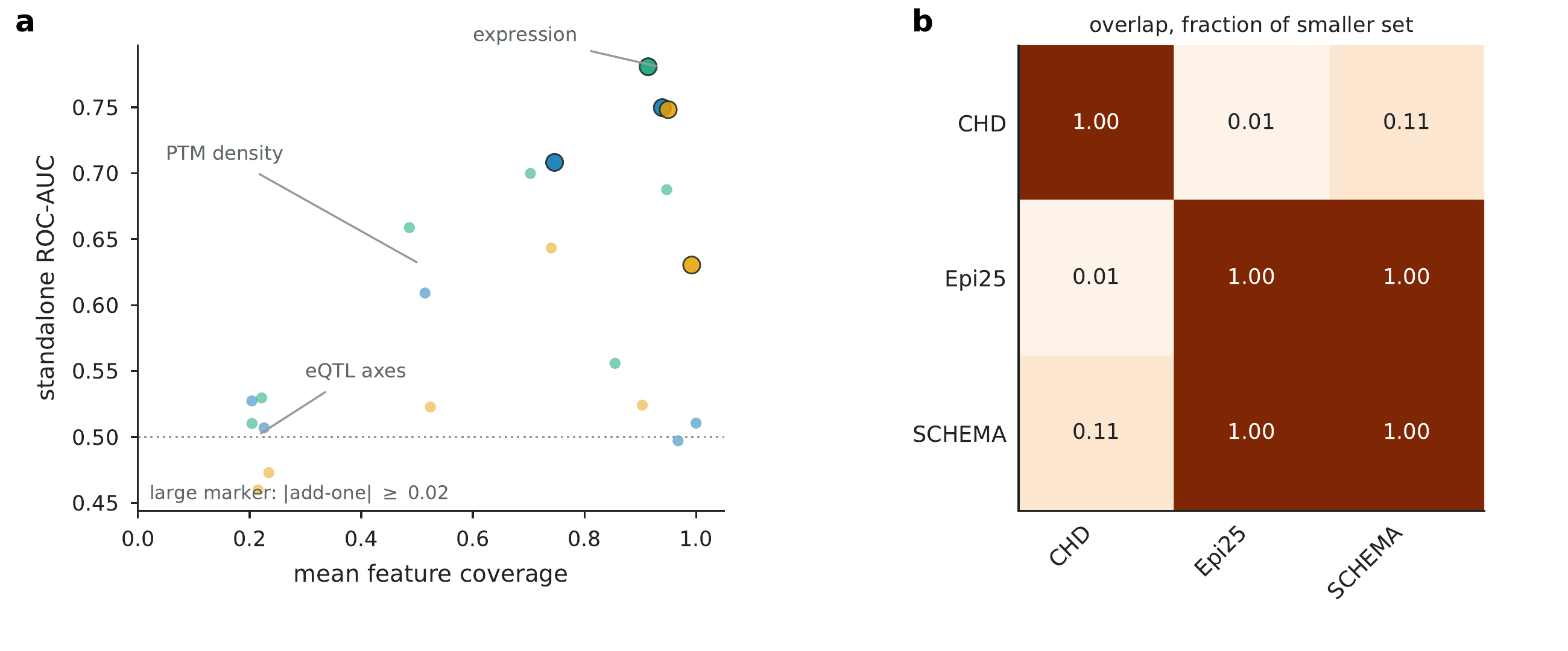

**Supplementary Figure S4. Diagnostics of the feature groups and overlap between disease labels.** **(a)** Each point is one feature group in one cohort (blue: CHD, orange: Epi25, green: SCHEMA) on the complete evaluated gene universe. X-axis denotes the mean coverage of the feature group, that is, the fraction of genes with a value, and Y-axis its standalone ROC-AUC. Large markers denote feature groups with an add-one ΔAUC over the baseline of at least 0.02 in absolute value, and the dotted line denotes chance. Values are given in **Supplementary Table S4**. **(b)** Overlap between the curated disease genes of the three cohorts, as a fraction of the smaller set.

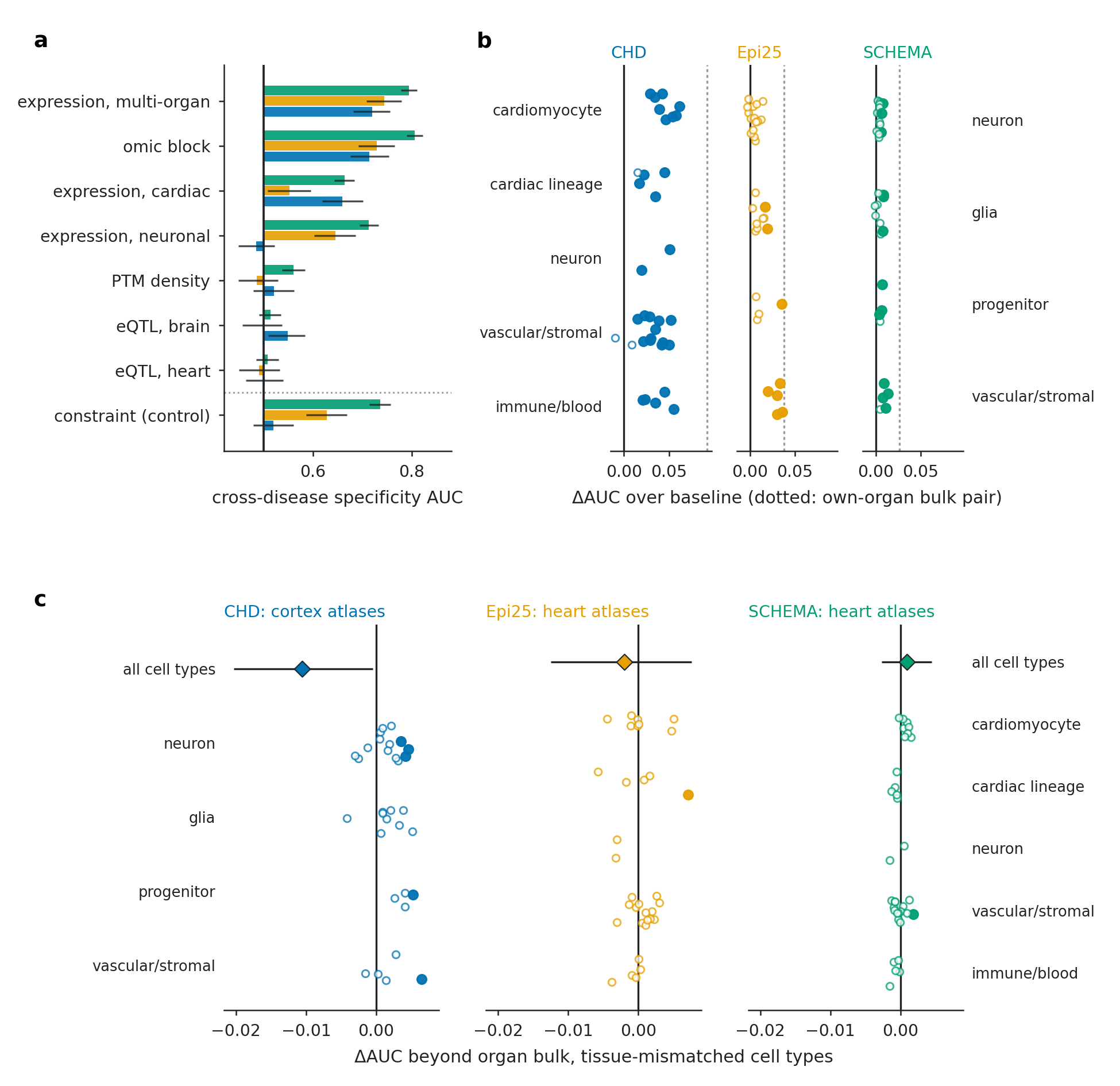

**Supplementary Figure S5. Cross-disease specificity and cell-type expression.** **(a)** ROC-AUC of each feature group among curated Mendelian disease genes (3,952, 3,871 and 4,157 genes for CHD, Epi25 and SCHEMA), using the cohort's disease genes as positives and genes associated with other diseases as negatives. The omic block includes all multi-omics features, and gene constraint is shown as a control. Values are given in **Supplementary Table S5**. **(b)** ΔAUC over the baseline of each disease-matched cell type (heart for CHD, cortex for Epi25 and SCHEMA), added individually. Dotted lines denote the ΔAUC of the disease organ's bulk expression (human and mouse) over the same baseline. **(c)** ΔAUC of tissue-mismatched cell types, added individually to the baseline and the multi-organ bulk expression. Diamonds denote all cell types added together. Filled points denote 95% confidence intervals excluding zero (2,000 bootstrap resamples).

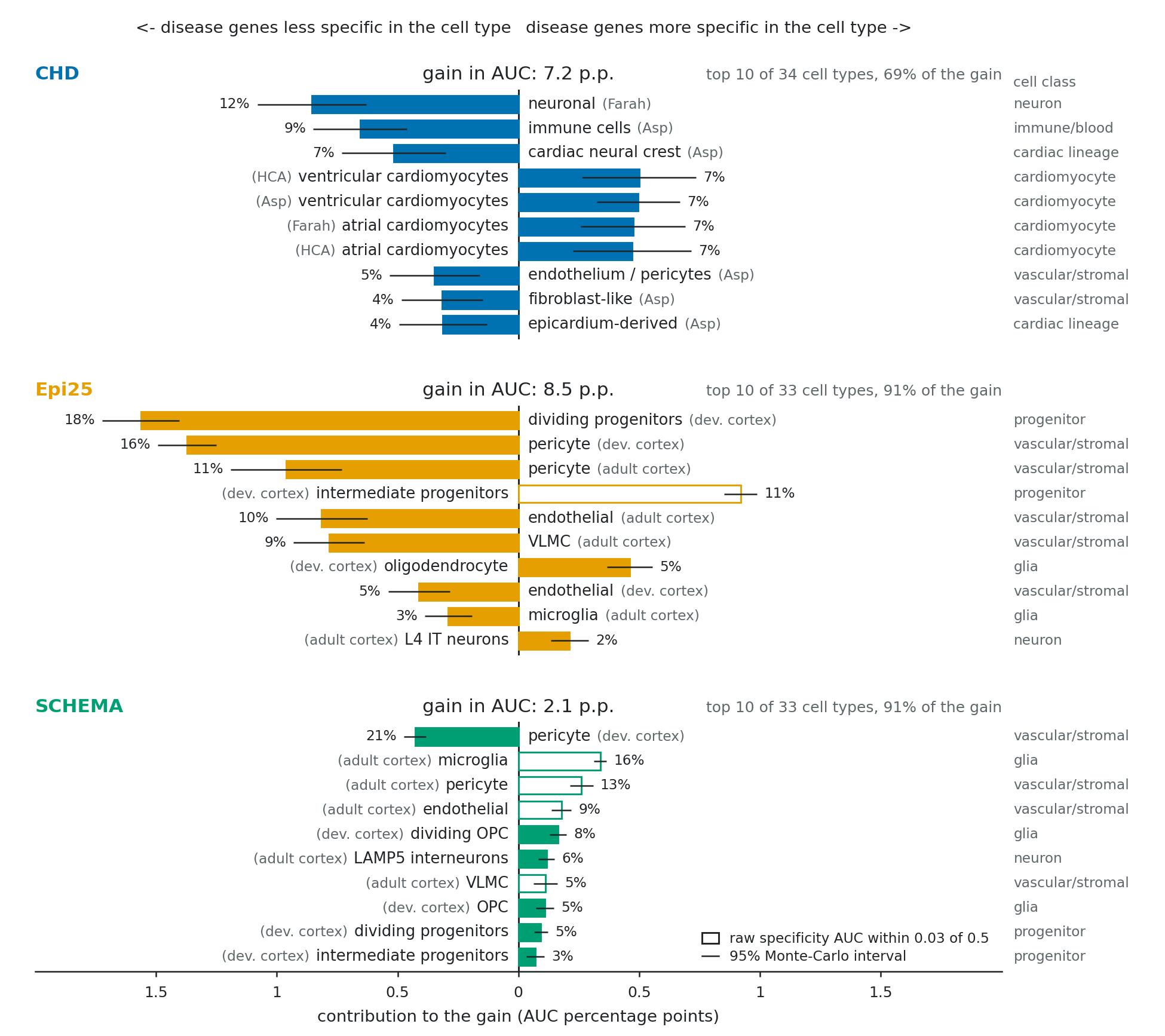

**Supplementary Figure S6. Cell-type contributions to the single-cell expression gain.** The ΔAUC of the single-cell profile over the baseline (Figure 4b) split over cell types (Monte-Carlo Shapley, 100 random orders), with the total gain in AUC percentage points shown above each panel. The ten largest contributors are shown for each cohort, with the source atlas in parentheses (Asp [1] and Farah [2]: developing heart, HCA: adult Heart Cell Atlas [3], dev. cortex and adult cortex: cortex atlases) and the cell class on the right. Bars to the right denote known genes more specific to that cell type than other genes, and bars to the left less specific. Hollow bars denote a specificity AUC within 0.03 of 0.5, and whiskers the 95% Monte-Carlo intervals.

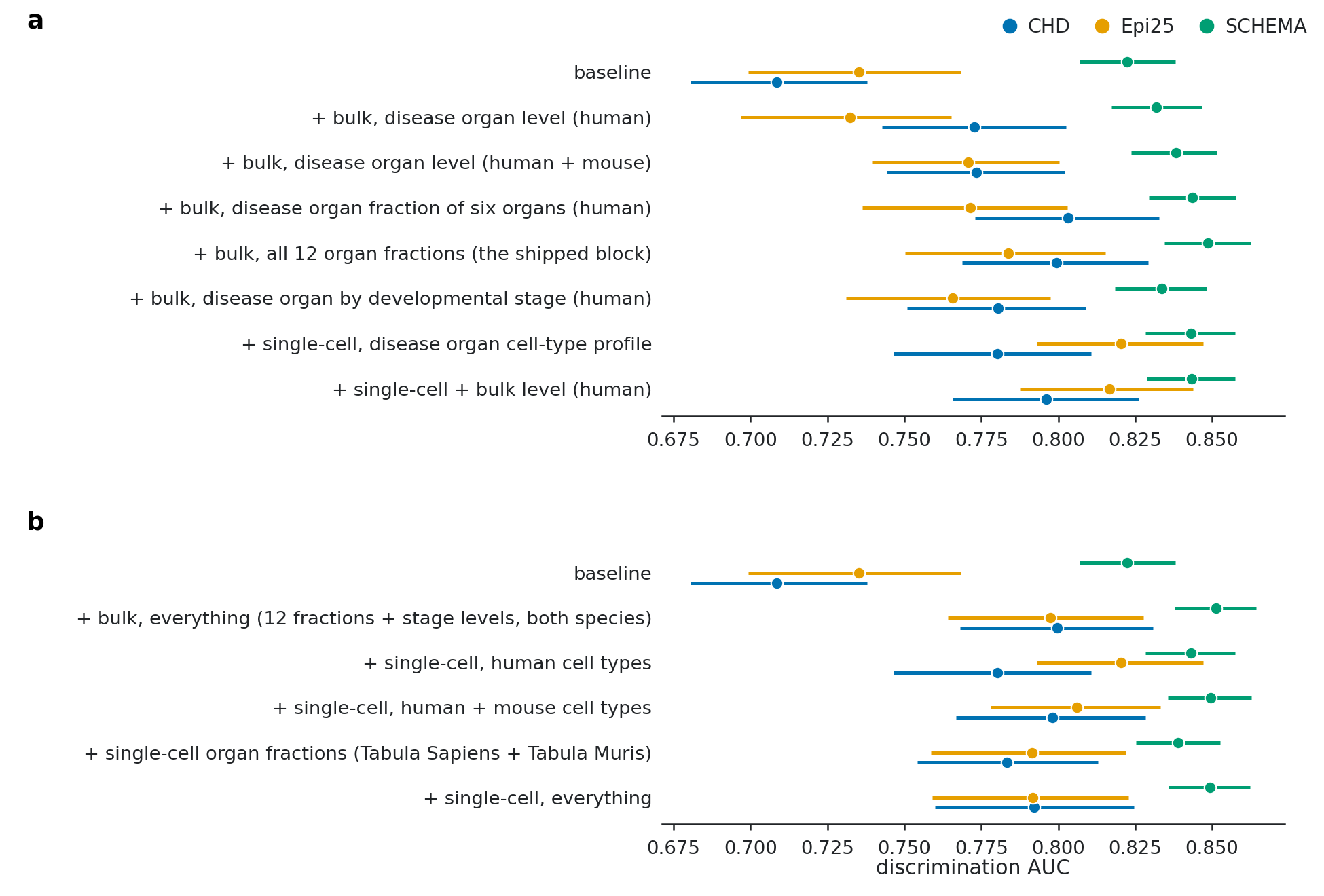

**Supplementary Figure S7. Contribution of bulk and single-cell expression.** Out-of-fold ROC-AUC with 95% confidence intervals for the cohort baseline (top row) and for the baseline plus each expression feature set (blue: CHD, orange: Epi25, green: SCHEMA). **(a)** Developmental bulk expression [4], from a single column for the disease organ (the bulk comparator in **Figure 4b**) to all organ fractions and developmental stages, compared with the single-cell cell-type profile. **(b)** Single-cell expression, from human cell types to organ-level fractions from Tabula Sapiens [5] and Tabula Muris [6], compared with all bulk features.

### Supplementary Tables

**Supplementary Table S1. ΔAUC of multi-omics features over three reference models.** Multi-omics features were added to gene constraint, to the cohort baseline (burden statistic, gene constraint and publication count) and to a model with the four published gene-prior scores fitted together (DOMINO, s_het_, pHaplo and GeVIR). Values denote ΔAUC [95% CI] on the complete gene universe. All scores are out of fold.

| **Cohort** | **Genes / positives** | **Over gene constraint** | **Over cohort baseline** | **Over fitted gene priors** |
| --- | --- | --- | --- | --- |
| CHD | 16,349 / 234 | +0.139 [+0.099, +0.181] | +0.089 [+0.054, +0.124] | +0.139 [+0.100, +0.179] |
| Epi25 | 15,956 / 200 | +0.091 [+0.056, +0.126] | +0.054 [+0.026, +0.086] | +0.083 [+0.048, +0.119] |
| SCHEMA | 18,569 / 710 | +0.034 [+0.022, +0.045] | +0.028 [+0.019, +0.038] | +0.031 [+0.020, +0.042] |

**Supplementary Table S2. Discrimination of the gene-prior scores and ranking models.** Values denote ROC-AUC [95% CI] for curated disease genes on the complete gene universe, with all models evaluated on the same cross-validation folds. AUC values should be compared within a cohort only, since gene universes and the number of positives differ between cohorts.

| **Score** | **CHD** | **Epi25** | **SCHEMA** |
| --- | --- | --- | --- |
| *Published gene-prior scores* | | | |
| DOMINO | 0.670 [0.631, 0.708] | 0.632 [0.591, 0.673] | 0.691 [0.670, 0.713] |
| s_het_ (GeneBayes) | 0.622 [0.584, 0.657] | 0.704 [0.670, 0.737] | 0.779 [0.762, 0.797] |
| pHaplo | 0.665 [0.628, 0.699] | 0.650 [0.614, 0.686] | 0.716 [0.697, 0.736] |
| GeVIR | 0.659 [0.625, 0.693] | 0.721 [0.685, 0.757] | 0.787 [0.770, 0.804] |
| All four, fitted | 0.646 [0.609, 0.684] | 0.681 [0.646, 0.717] | 0.808 [0.792, 0.824] |
| *Reference models* | | | |
| Publication count | 0.663 [0.632, 0.695] | 0.617 [0.586, 0.647] | 0.691 [0.675, 0.707] |
| Gene constraint | 0.634 [0.600, 0.667] | 0.678 [0.638, 0.716] | 0.809 [0.791, 0.826] |
| Cohort baseline (burden + constraint + publication count) | 0.708 [0.680, 0.738] | 0.735 [0.699, 0.768] | 0.822 [0.807, 0.838] |
| *With multi-omics features* | | | |
| Gene constraint + multi-omics | 0.774 [0.738, 0.810] | 0.769 [0.736, 0.801] | 0.843 [0.829, 0.857] |
| Baseline + multi-omics (re-rank) | 0.797 [0.765, 0.827] | 0.790 [0.758, 0.821] | 0.851 [0.837, 0.864] |

**Supplementary Table S3. Contribution of each feature group by removal from the full model.** Values denote the decrease in AUC [95% CI] when one feature group is removed from the full model (baseline plus multi-omics features), on the complete gene universe. CVPC: cardiovascular progenitor cells.

| **Feature group removed** | **CHD** | **Epi25** | **SCHEMA** |
| --- | --- | --- | --- |
| Multi-organ bulk expression | +0.024 [0.000, +0.047] | +0.026 [+0.002, +0.051] | +0.019 [+0.011, +0.026] |
| eQTL, heart | +0.012 [+0.003, +0.022] | +0.001 [-0.004, +0.006] | -0.001 [-0.003, +0.001] |
| Cardiac expression | +0.001 [-0.013, +0.014] | +0.009 [-0.004, +0.021] | +0.002 [-0.001, +0.005] |
| eQTL, brain | -0.006 [-0.011, -0.001] | +0.003 [-0.003, +0.009] | -0.001 [-0.002, +0.001] |
| Neuronal expression | +0.002 [-0.003, +0.007] | +0.001 [-0.005, +0.008] | -0.001 [-0.002, +0.001] |
| PTM density | +0.001 [-0.004, +0.007] | 0.000 [-0.002, +0.003] | 0.000 [-0.001, 0.000] |
| eQTL, CVPC | 0.000 [0.000, 0.000] | +0.001 [-0.002, +0.003] | 0.000 [-0.001, 0.000] |

**Supplementary Table S4. Coverage and discrimination of each feature group.** Coverage denotes the mean fraction of genes with a value, standalone AUC the ROC-AUC of the feature group alone, and add-one ΔAUC the change in AUC when the feature group is added to the cohort baseline, on the complete gene universe. CVPC: cardiovascular progenitor cells.

| **Feature group** | **Varying / total columns** | **Coverage** | **Standalone AUC** | **Add-one ΔAUC** |
| --- | --- | --- | --- | --- |
| *CHD* | | | | |
| Multi-organ bulk expression | 12 / 12 | 0.94 | 0.750 | +0.091 |
| Cardiac expression | 6 / 6 | 0.75 | 0.708 | +0.079 |
| Neuronal expression | 2 / 2 | 0.97 | 0.497 | -0.004 |
| eQTL, heart | 5 / 5 | 0.23 | 0.507 | -0.002 |
| eQTL, brain | 5 / 5 | 0.20 | 0.527 | -0.006 |
| eQTL, CVPC | 1 / 1 | 1.00 | 0.511 | +0.001 |
| PTM density | 1 / 3 | 0.51 | 0.609 | +0.008 |
| *Epi25* | | | | |
| Multi-organ bulk expression | 12 / 12 | 0.95 | 0.748 | +0.049 |
| Cardiac expression | 6 / 6 | 0.74 | 0.643 | +0.013 |
| Neuronal expression | 2 / 2 | 0.99 | 0.630 | +0.020 |
| eQTL, heart | 5 / 5 | 0.23 | 0.473 | -0.010 |
| eQTL, brain | 5 / 5 | 0.22 | 0.460 | +0.010 |
| eQTL, CVPC | 1 / 1 | 0.90 | 0.524 | +0.003 |
| PTM density | 1 / 3 | 0.52 | 0.523 | -0.002 |
| *SCHEMA* | | | | |
| Multi-organ bulk expression | 12 / 12 | 0.91 | 0.781 | +0.026 |
| Cardiac expression | 6 / 6 | 0.70 | 0.700 | +0.007 |
| Neuronal expression | 2 / 2 | 0.95 | 0.688 | +0.006 |
| eQTL, heart | 5 / 5 | 0.22 | 0.530 | -0.002 |
| eQTL, brain | 5 / 5 | 0.20 | 0.510 | -0.001 |
| eQTL, CVPC | 1 / 1 | 0.85 | 0.556 | +0.003 |
| PTM density | 1 / 3 | 0.49 | 0.659 | 0.000 |

**Supplementary Table S5. Cross-disease specificity of each feature group.** Values denote ROC-AUC [95% CI] among curated Mendelian disease genes, using the cohort's disease genes as positives and genes associated with other diseases as negatives. Gene constraint is included as a control. CVPC: cardiovascular progenitor cells.

| **Feature group** | **CHD (n=3,952)** | **Epi25 (n=3,871)** | **SCHEMA (n=4,157)** |
| --- | --- | --- | --- |
| All multi-omics features | 0.714 [0.676, 0.753] | 0.729 [0.692, 0.765] | 0.806 [0.790, 0.823] |
| Multi-organ bulk expression | 0.720 [0.681, 0.756] | 0.744 [0.708, 0.779] | 0.794 [0.778, 0.810] |
| Cardiac expression | 0.659 [0.619, 0.701] | 0.553 [0.509, 0.596] | 0.664 [0.643, 0.684] |
| Neuronal expression | 0.485 [0.449, 0.523] | 0.645 [0.602, 0.686] | 0.713 [0.694, 0.733] |
| eQTL, heart | 0.502 [0.465, 0.539] | 0.491 [0.451, 0.533] | 0.508 [0.486, 0.530] |
| eQTL, brain | 0.548 [0.509, 0.584] | 0.498 [0.458, 0.538] | 0.514 [0.491, 0.535] |
| eQTL, CVPC | 0.527 [0.490, 0.566] | 0.524 [0.486, 0.563] | 0.474 [0.451, 0.499] |
| PTM density | 0.521 [0.479, 0.562] | 0.487 [0.449, 0.529] | 0.561 [0.537, 0.584] |
| Gene constraint (control) | 0.520 [0.479, 0.561] | 0.628 [0.586, 0.669] | 0.736 [0.714, 0.757] |

**Supplementary Table S6. Discrimination of disease genes by organ expression.** Values denote ROC-AUC [95% CI] of each organ's developmental bulk expression [4] among curated Mendelian disease genes. The disease-matched organ (bold) ranked first in all three cohorts. In Epi25, the margin over the second organ (testis) was the least certain (P=0.029, and P=0.094 conditional on gene constraint).

| **Organ** | **CHD** | **Epi25** | **SCHEMA** |
| --- | --- | --- | --- |
| Brain | 0.564 [0.525, 0.601] | **0.720 [0.681, 0.758]** | **0.773 [0.756, 0.790]** |
| Heart | **0.711 [0.672, 0.748]** | 0.527 [0.486, 0.570] | 0.595 [0.574, 0.616] |
| Kidney | 0.588 [0.548, 0.627] | 0.577 [0.535, 0.617] | 0.611 [0.590, 0.632] |
| Liver | 0.603 [0.571, 0.637] | 0.556 [0.516, 0.598] | 0.645 [0.624, 0.666] |
| Ovary | 0.567 [0.529, 0.604] | 0.616 [0.575, 0.655] | 0.580 [0.560, 0.602] |
| Testis | 0.513 [0.474, 0.550] | 0.668 [0.628, 0.705] | 0.619 [0.599, 0.640] |

**Supplementary Table S7. Recovery of known disease genes by each ranking method.** Number (%) of curated disease genes in the top decile of each cohort's gene universe, as shown in **Figure 4a**, followed by further ranking summaries for the cohort baseline and the re-rank model (baseline plus multi-omics features). The paired difference is given with its 95% CI.

| **Summary** | **CHD** | **Epi25** | **SCHEMA** |
| --- | --- | --- | --- |
| *Known genes in the top decile (Figure 4a)* | | | |
| Genes in the top decile | 1,635 | 1,596 | 1,857 |
| Fitted gene-prior scores | 72 (30.8%) | 59 (29.5%) | 336 (47.3%) |
| Cohort baseline | 61 (26.1%) | 81 (40.5%) | 359 (50.6%) |
| Baseline + multi-omics (re-rank) | 115 (49.1%) | 95 (47.5%) | 380 (53.5%) |
| Re-rank minus baseline, percentage points [95% CI] | +23.1 [+15.2, +30.5] | +7.0 [+0.5, +13.9] | +3.0 [+0.9, +5.8] |
| *Other ranking summaries (baseline / re-rank)* | | | |
| Genes ranked to recover half of the known genes | 4,236 / 1,755 | 3,245 / 2,074 | 1,826 / 1,599 |
| Known genes among the top 100 genes | 5% / 39% | 23% / 27% | 52% / 54% |
| Average precision | 0.035 / 0.180 | 0.073 / 0.112 | 0.243 / 0.258 |
| AUC of the multi-omics features alone | 0.778 [0.743, 0.811] | 0.766 [0.734, 0.797] | 0.799 [0.785, 0.814] |

### Supplementary References

[1]	Asp M, Giacomello S, Larsson L, Wu C, Fürth D, Qian X, et al. A spatiotemporal organ-wide gene expression and cell atlas of the developing human heart. Cell 2019;179(7):1647-1660. doi:10.1016/j.cell.2019.11.025.

[2]	Farah EN, Hu RK, Kern C, Zhang Q, et al. Spatially organized cellular communities form the developing human heart. Nature 2024;627:854-864. doi:10.1038/s41586-024-07171-z.

[3]	Litviňuková M, Talavera-López C, Maatz H, Reichart D, Worth CL, Lindberg EL, et al. Cells of the adult human heart. Nature 2020;588:466-472. doi:10.1038/s41586-020-2797-4.

[4]	Cardoso-Moreira M, Halbert J, Valloton D, Velten B, Chen C, Shao Y, et al. Gene expression across mammalian organ development. Nature 2019;571(7766):505-509. doi:10.1038/s41586-019-1338-5.

[5]	Tabula Sapiens Consortium. The Tabula Sapiens: a multiple-organ, single-cell transcriptomic atlas of humans. Science 2022;376(6594):eabl4896. doi:10.1126/science.abl4896.

[6]	Tabula Muris Consortium. Single-cell transcriptomics of 20 mouse organs creates a Tabula Muris. Nature 2018;562(7727):367-372. doi:10.1038/s41586-018-0590-4.
